# Ice gliding diatoms establish record-low temperature limit for motility in a eukaryotic cell

**DOI:** 10.1101/2024.11.18.624199

**Authors:** Qing Zhang, Hope T. Leng, Hongquan Li, Kevin R. Arrigo, Manu Prakash

## Abstract

Despite periods of permanent darkness and extensive ice coverage in polar environments, photosynthetic ice diatoms display a remarkable capability of living inside the ice matrix. How these organisms navigate such hostile conditions with limited light and extreme cold remains unknown. Using a custom sub-zero temperature microscope during an Arctic expedition, we present the discovery of motility at record-low temperatures in a Eukaryotic cell. By characterizing the gliding motility of several ice diatom species, collected from ice cores in the Chukchi Sea, we record that they retain motility at temperatures as low as –15 °C. Remarkably, ice diatoms can glide on ice substrates, a capability absent in temperate diatoms of the same genus. This unique ability arises from adaptations in extracellular mucilage that allow ice diatoms to adhere to ice, essential for gliding. Even on glass substrates where both cell types retain motility at freezing temperatures, ice diatoms move an order of magnitude faster, with their optimal motility shifting toward colder temperatures. Combining field and laboratory experiments with thermo-hydrodynamic modeling, we reveal adaptive strategies that enable gliding motility at extreme cold temperatures. These strategies involve increasing internal energy efficiency with minimal changes in heat capacity and activation enthalpy, and reducing external dissipation by minimizing the temperature sensitivity of mucilage viscosity. The discovery of diatoms’ ice gliding motility opens new routes for understanding their survival within a harsh ecological niche and their migratory responses to rapid environmental changes. Our work highlights the robust adaptability of ice diatoms in one of Earth’s most extreme settings.

## Introduction

Ice diatoms are important components of algal communities in polar regions, serving as the major primary producers prior to the spring phytoplankton bloom^1–3^. These microorganisms exhibit extraordinary adaptations to the extreme conditions within sea ice^4–11^. Polar ice caps envelop up to 13% of the Earth’s ocean surface and are characterized by internal porous structures filled with brine channels, creating a unique habitat with subfreezing temperatures and highly fluctuating salinity^12–14^. Despite the harsh conditions, ice diatoms have developed specialized adaptations to remain active, including increased efficiency of metabolism and fluidity of lipid compositions^6,15–18^. While some biochemical adaptations that enable ice diatoms to thrive under extreme cold are known, their locomotive strategies remain largely unexplored.

Ice diatoms’ behavioral adaptations critically influence their survival and interaction with the icy habitat^19,20^. They are believed to ‘select’ specific depths within the ice core that offer optimal light, nutrients, and salinity, enhancing their survival prospects^21–26^. Furthermore, ice diatoms secrete substances such as ice-binding proteins (IBPs) and extracellular polymeric substances (EPS)^27–29^. These substances, while protecting the diatoms from cold and balancing osmotic stress^16,30^, also alter the process of ice crystallization^31–34^ and impact polar ocean carbon cycling^30,35,36^. The dual ability of ice diatoms to relocate within the ice for survival and to reshape their habitat highlights the importance of understanding their motility mechanism. By migrating within brine channel networks in sea ice, which becomes permeable at temperatures above −5 °C^12^, diatoms can spread and affect polar ecosystems, including initiating new under-ice blooms^37^. It has been hypothesized that the diatoms living in brine channels might seed under-ice blooms due to increased ice porosity during warming spring^38^. As polar regions undergo rapid changes in climate, it is urgent to understand the adaptive physiology of these unique species living in ice and how they might evolve with rising temperatures^32,39,40^.

Despite the critical role of migration ability, there remains a significant gap in our understanding of ice diatoms’ motility in their native icy environments. Indirect empirical evidence, such as changing ice thickness leading to the relocation of algal colonies at the ice bottom, has provided estimates of diatom movement, suggesting a minimum velocity of approximately 1.5 cm per day^41^. Discrepancies between observed cell densities and photosynthetic measurements in ice, unexplained by local growth rates alone, further suggest active migration of cells within the ice^42^. Additionally, field experimental manipulations of snow depth, which influence light penetration, have demonstrated that ice diatoms can strategically reposition themselves within the ice column in a few days^43^. These behavioral adjustments, believed to be adaptive responses to shifting light conditions, suggest an active engagement with their environment rather than passive diffusion^43^. Following laboratory observations of an ice diatom species, *Cylindrotheca closterium*, reveal that it can move on the surface of a culture flask at 0 °C^43^. However, translating this finding to the context of natural ice environments introduces uncertainties, because the physical and chemical properties of laboratory surfaces differ significantly from those of sea ice. The lack of direct observations within the actual ice context leaves critical questions about ice diatom motility, including the specific mechanisms they employ and their responses to environmental changes like temperature fluctuations.

In this study, we present direct cellular observations of ice diatoms demonstrating unique ice gliding ability across their natural habitat in sea ice. This ability reflects their adaptations to two critical factors: icy substrates and cold temperatures. First, we show that only ice diatoms can move on ice; temperate diatoms immediately lose motility upon contact due to lack of adhesion to ice. Second, we find that ice diatoms maintain enhanced motility at freezing temperatures. On glass substrates where both types can move, ice diatoms exhibit gliding speeds an order of magnitude faster than their temperate counterparts. To understand the mechanisms behind this superior cold motility, we investigate temperature-dependent factors governing gliding speed. By introducing traction force microscopy (TFM) tailored for diatoms, we map the forces applied and experienced by gliding diatoms, revealing distinct interactions with the substrate. Based on experimental insights, we develop a thermo-hydrodynamic model incorporating primary temperature-dependent factors for gliding motility, revealing strategies ice diatoms have evolved to maintain efficient movement under extreme cold conditions. Furthermore, we investigate population motility of ice diatoms and reveal temperature-dependent patterns characterized by Gamma and Chi-squared distributions. As temperature increases, ice diatoms diversify gliding speeds, indicating adaptive behaviors that enhance ecological success under changing conditions.

## Results

### Unique gliding motility of Arctic diatoms on ice

To investigate the dynamic behavior of ice diatoms within sea ice, we conduct a 45-day Arctic expedition aboard the R/V Sikuliaq, collecting ice cores from the first-year sea ice in the Chukchi Sea (70.3° - 71.6° N, 161.0° - 165.8° W) from 12 ice stations throughout the summer season in 2023 (June 15 to July 30) (Fig. 1a and Table S1). These cores feature biota layers embedded in the ice, which appear brown and with reduced transparency at specific depths, as shown in Fig. 1a (ice station 21 at 71.3° N, 164.6° W).

**Figure 1.**
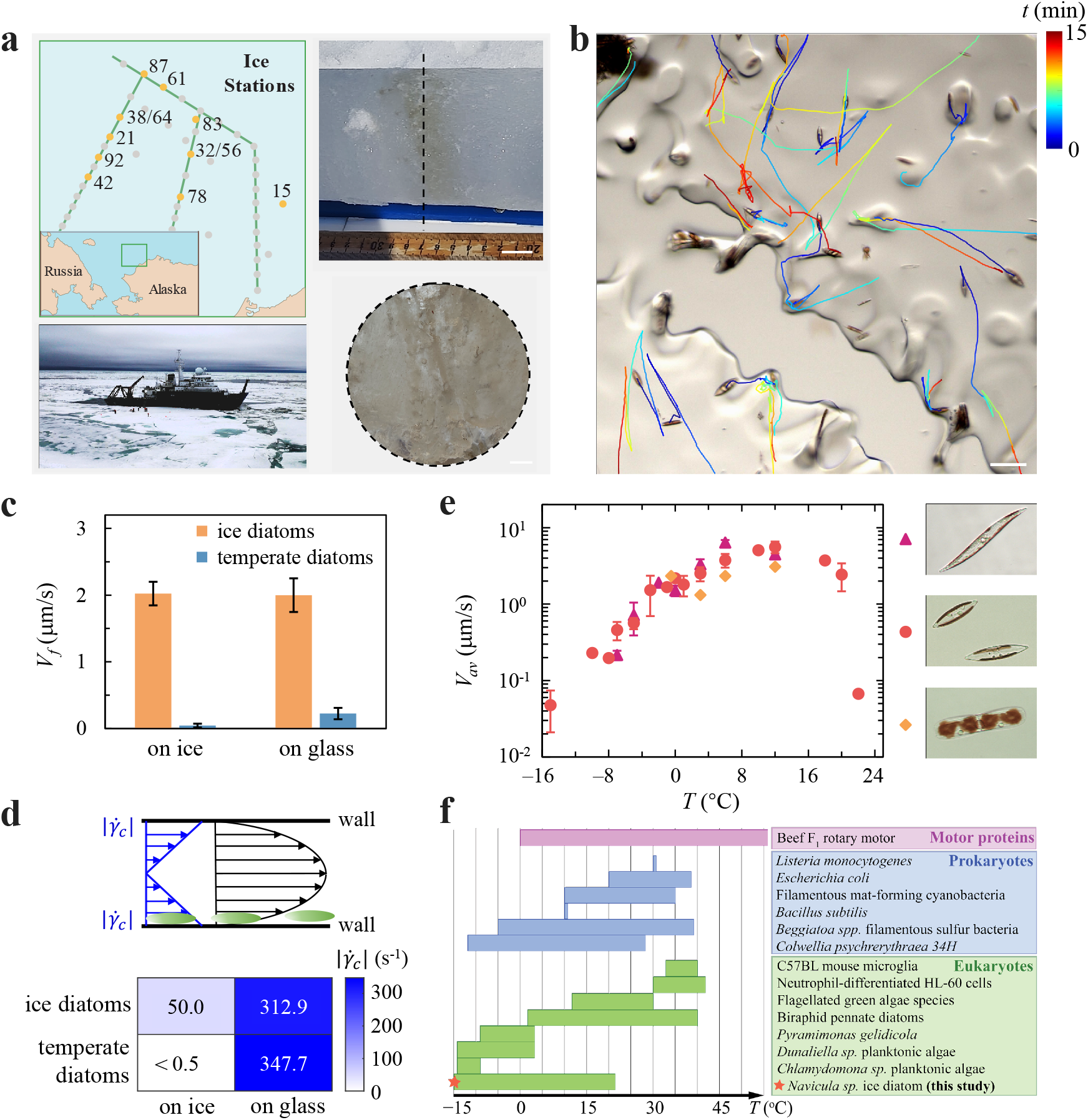
Gliding motility of ice diatoms. (**a**) Left panel: Expedition route and ice stations in the Arctic. Upper right panel: Biotic layer observed in an Arctic ice core from the Chukchi Sea. Scale bars: 20 mm. Lower right panel: Cross-sectional view of the ice core showing the region with the highest diatom density. Scale bars: 2 mm. (**b**) Time-lapse trajectories of ice diatoms gliding directly on an ice surface, imaged using sub-zero temperature controlled microscopy. Scale bar: 100 *μ*m. (**c**) Comparison of the gliding motility of ice diatoms (*Navicula sp*. and *Pluerosigma sp*., blue) and temperate diatoms (*Navicula arenaria var. rostellata, Navicula sp*. and *Pluerosigma sp*., orange) reveals that ice diatoms retain the ability to glide at 0 °C. In contrast, temperate diatoms can still glide on glass substrates at this temperature with a speed an order of magnitude lower but completely lose gliding ability on ice. This indicates that both substrate and ambient temperature influence gliding motility. (**d**) Schematic of direct adhesion strength measurements for ice and temperate diatoms on ice and glass surfaces using shear generated by Poiseuille flow (Upper panel). On glass substrates, both types of diatoms adhere, whereas on ice substrates, only ice diatoms maintain adhesion (Lower panel). (**e**) Average gliding speed as a function of temperature for three species of ice diatoms collected in the field: 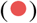 *Navicula sp*., 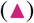 *Pluerosigma sp*., 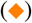 *Entomoneis sp*.. (**f**) Temperature dependence of cellular motility and transport across the tree of life, illustrating that ice diatoms approach the lowest temperature limits of motility^44–55^.

At each ice station, we systematically image the diatoms under high magnification (10x, 20x, 40x) to understand the biodiversity of ice diatoms trapped in ice cores. Although exact species have not been identified, we predominantly find pennate diatoms such as *Navicula sp*., *Pleurosigma sp*., *Nitzschia frigida, Entomoneis sp*., *Pseudo-nitzschia* and *Fragilariopsis sp*, identified based on their morphological characteristics (Fig. S1). Our microscopic observations reveal that the diatoms presenting in the ice core can also be found in the water column.

While the presence of diatoms in ice cores has been reported for decades^2,56^, no cellular-scale observations of their behavior in ice exist. Ice is a scattering media, making direct observation challenging. We circumnavigate this issue by extracting diatoms from ice cores and re-dispersing them on thin ice film surfaces or within micro-ice channels (see Methods for details). These preparations are then examined under high magnification (4x, 10x) at desired temperatures using our customized temperature-controlled microscope (see Methods for details). Remarkably, we find that the Arctic pennate diatom species can actively glide on ice surfaces and within ice channels, as shown in Figs. 1b and S2, and Movies S1 and S2. Although salinity in natural brine channels varies with temperature^26^, the salinity in our experiments remains effectively constant (approximately 36 ppt), because the timescale of diatoms’ motility is much shorter than that of the ice film growth during our observations.

To illustrate the uniqueness of ice gliding motility, we compare the ice diatom *Navicula sp*. with various temperate species (*Navicula sp*., *Navicula arenaria var. rostellata*, and *Pleurosigma sp*.) gliding on both glass and ice substrates. On glass surfaces at 0 °C, although both ice and temperate diatoms exhibit motility, the ice diatoms move significantly faster–nearly an order of magnitude greater than the temperate counterparts, as shown in Fig. 1c and Movie S3. This enhanced motility reveals remarkable cold-adapted motility. Interestingly, on ice surfaces, temperate diatoms completely lose their ability to glide; any movement observed is limited to passive drifting due to uneven ice surfaces or self-interactions in aggregations (Fig. 1c and Movie S3). In contrast, ice diatoms maintain consistent gliding speeds comparable to those observed on glass, as shown in Fig. 1c. This stark contrast indicates the unique ability of ice diatoms to navigate on and through ice, suggesting that they have evolved specialized physiological traits to interact effectively with their icy habitat. These comparisons indicate two exceptional aspects regarding the adaptations of ice diatoms’ motility: i) the unique ability to interact with an icy substrate; ii) the motile resilience to cold temperatures.

Gliding motility in diatoms relies on typical interactions with the substrate. Diatoms are known to secrete mucilage to adhere to surfaces, which facilitates their motility^57–60^. We quantify the adhesion strength of this mucilage by measuring the shear rate required to detach diatoms from surfaces in a fluidic channel (detailed in the Methods section). Our measurements show that ice diatoms maintain considerable adhesion levels on both ice and glass surfaces, with shear strain tolerances of 50.0 s^−1^ and 312.9 s^−1^, respectively. In contrast, their temperate counterparts fail to adhere to ice at the minimal shear strain of 0.5 s^−1^ (the lowest we can apply) but adhere well to glass with a strength of 347.7 s^−1^, as shown in Fig. 1d. These behaviors align with their respective motility on different substrates (Fig. 1e). The alignment between the mapping of motility and adhesion strength across different substrates suggests that the mucilage secreted by ice diatoms may include specific ice binding capabilities that enable attachment to ice and contribute to the unique gliding ability on ice (see Discussion).

Temperature is a fundamental state variable which influences nearly all cellular processes, including those governing motility^61–64^. Despite its significance, cellular motility under extreme cold conditions has not been extensively studied^45,52,53,65,66^. To assess the impact of temperature variations on ice diatoms’ motility, we quantify the average moving speed of three prevalent wild ice diatoms in the Arctic, *i*.*e*., *Navicula sp*., *Pleurosigma sp*., and *Entomoneis sp*.. The majority of data are collected within 24-48 hrs from the moment of sampling using the customized temperature-controlled microscope aboard the R/V Sikuliaq. Remarkably, all three species remain motile at freezing or subfreezing temperatures, and *Navicula sp*. is able to move down to −15 °C. Moreover, these species show a general trend where their average gliding speed increases with rising temperatures, reaching a plateau at around 12 °C (defined as optimal temperature), and then declining sharply to near zero at approximately 22 °C (defined as maximum temperature), as shown in Fig. 1e. In contrast, temperate diatoms in the same genera (*Pleurosigma sp*. and *Navicula sp*.) exhibit an optimal motility temperature around 30 °C and a maximum deactivating temperature near 35 - 40 °C (See Supplementary Information and Fig. S3 for details). When exposed to colder temperatures, these temperate diatoms almost cease their motion at approximately −1 °C. The systematic shift of temperature response curves toward lower temperatures corresponds with the enhanced motility at freezing temperatures for ice diatoms (Fig. 1c) and highlights notable adaptations to cold. We explore the underlying mechanisms that counteract the inhibitory effects of extreme cold on gliding processes in the following section.

Having established the unique ability of ice diatoms to glide on ice substrates and their enhanced motility at cold temperatures, we wonder how this compares to the broader context of cellular motility across the tree of life. By surveying published results on the impact of temperature on motility in single cells and the related activity of motor proteins^44–55,67–70^, we create a comprehensive dataset that includes the lowest temperature experiments performed to date, as shown in Fig. 1f (up to 50 °C) and Fig. S4. Notably, we find that ice diatoms exhibit gliding motility at the lowest recorded temperature limits in a Eukaryotic cell. Their adaptation across a broad range of temperatures provides a possible explanation for why they form dominant communities in ice cores (Fig S1)^1,9,56^.

### Strategies for cold-adapted motility in *Navicula sp*

To understand the strategies enabling ice diatoms to adapt their motility to freezing temperatures, we concentrate our investigations on an Arctic species, *Navicula sp*., which we isolate and fortunately culture from an ice core at Ice Station 87 (71.7° N, 164.0° W) in the Chukchi Sea (see Methods for details). Our observations of similar gliding speeds at freezing temperatures on both ice and glass substrates (Fig. 1d) suggest that the mechanisms controlling their gliding speeds are independent of the type of substrates. Therefore, we choose flat glass substrates for our experiments to explore their gliding mechanisms and to dissect the temperature-dependent motility.

Gliding motility is a fascinating mode of transport utilized across the tree of life, both in prokaryotic and eukaryotic cells^71–73^. Although many studies have investigated the mechanisms of motility in pennate diatoms, a quantitative understanding of the key factors governing gliding speeds remains limited^57,74–76^. Gliding motility in temperate diatoms is powered by internal motor proteins, specifically myosin interacting with actin cables along the cell^77–79^. This process involves the secretion of adhesive mucilage threads that bind to the substrate and couple internally to the motile motors^60^. The myosin motors pull the mucilage threads toward the rear, using friction with the substrate to propel the diatom forward, known as mucilage-thread-myosin machinery^59,60,80,81^, as schematically shown in Fig. 2a. Employing similar experimental approaches as in temperate diatoms, we investigate the internal mechanisms driving the motility of ice diatoms.

**Figure 2.**
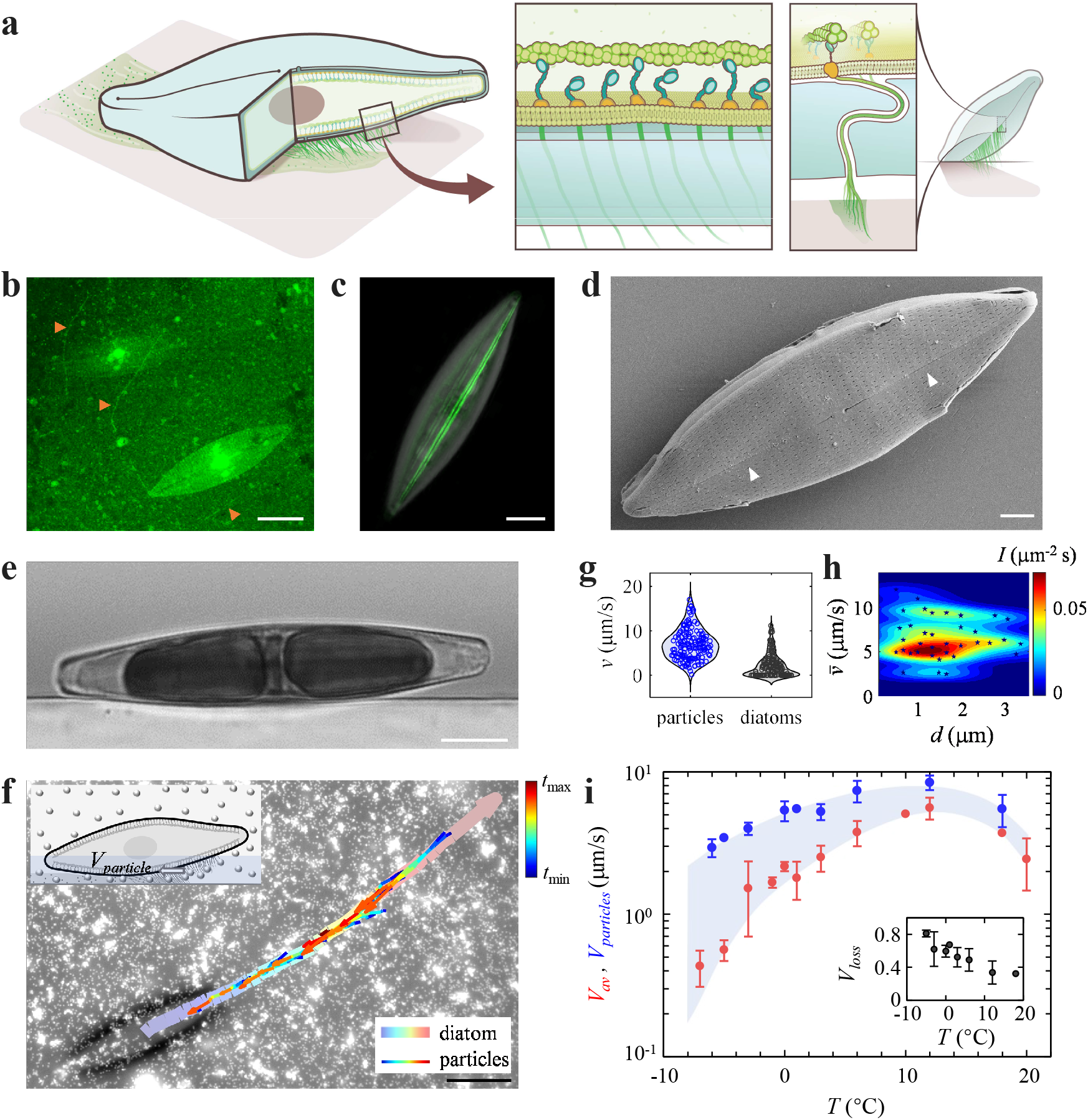
Physical factors governing the gliding speed of ice diatoms. (**a**) Schematic illustrates diatom gliding motility driven by acto-myosin and mucilage thread-based latching machinery. (**b**) Mucilage traces left by ice diatoms, *Navicula sp*., labeled with Wheat Germ Agglutinin (WGA), indicate movement paths (orange arrows). Scale bar: 20 *μ*m. (**c**) Three-dimensional visualization of actin cables (green) within *Navicula sp*., illustrates cytoskeletal structures crucial for gliding motility. Scale bar: 10 *μ*m. (**d**) Scanning Electron Micrograph of *Navicula sp*. reveals thin slits (raphes) on the diatom frustules (white arrows). Scale bar: 3 *μ*m. (**e**) Side view of a gliding ice diatom on a glass substrate shows a tilt resulting from torque competition between active and passive drag forces. Scale bar: 10 *μ*m. (**f**) A directional flux beneath *Navicula sp*. via raphe grooves is revealed using fluorescent polystyrene beads (0.5 *μ*m), indicating that tracer particles latched to the ventral raphe move opposite to diatom motion. Thick curve shows the diatom’s trajectory; thin curves show tracer particles’ trajectories. Inset: Schematic illustrates tracer particles latched to mucilage threads that are moving along the ventral raphe. The tracer particle movement reflects internal acto-myosin kinetics. Scale bar: 20 *μ*m. (**g**) Distribution of gliding velocities for diatoms and speeds of latched tracer particles, *v*, in the ventral region. (**h**) Average speed of tracer particle clusters attached to mucilage threads is independent of their size (projection diameter, *d*). Color map indicates intensity of kernel density estimation, *I*. (**i**) Comparison of temperature-dependent motility of ice diatoms absent of tracer particles, *V*_*av*_ 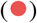 and mucilage latched tracer particle velocity, *V*_*particles*_ 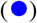 reveals significant reduction from velocity generated by internal machinery to final gliding velocity. Inset: Normalized velocity loss, *V*_*loss*_ = (*V*_*particles*_*− V*_*av*_) */V*_*particles*_, demonstrates the temperature-dependent impact of external boundary conditions.

The Arctic *Navicula sp*. is kept at 3 °C with a 12-hour day/night cycle. We examine their mucilage secretion by allowing these diatoms to glide on a glass surface for 12 hours and staining the substrate with Wheat Germ Agglutinin (see Methods for details). Fine mucilage residues left on the substrate are directly visualized, as shown in Fig. 2b. Additionally, we establish the actin cytoskeleton architecture in the ice diatom by labeling their actin filaments with phalloidin (see Methods for details). Two actin cables running parallel to the long axis are co-located within the raphe region (Fig. 2c). The raphe structure in ice diatoms is revealed in scanning electron microscopy (SEM), as shown in Fig. 2d. These observations demonstrate that ice diatoms utilize a gliding mechanism similar to that of temperate species^60^.

While powered by internal motor proteins, moving diatoms strongly interact with the substrate and the surrounding hydrodynamic environment. Side-view imaging of live gliding Arctic *Navicula sp*. show that these diatoms always tend to tilt during their movement, as shown in Fig. 2e and Movie S4. The tilted angles observed in gliding ice diatoms correlate well to previous observations on temperate diatom, *Craticula cuspidata*^78^. The observed tilt angles in both ice and temperate diatoms reflect the interplay between the driving torque from the mucilage-thread-myosin machinery and counteracting hydrodynamic torques (see Discussion).

Gliding motility of diatoms is governed by complex processes involving internal and environmental factors. To explore how temperature impacts these processes, we categorize the factors governing gliding speed into two temperature-dependent regimes: i) internal interactions, which encompass all temperature-modulated cellular dynamics such as the driving forces generated by the mucilage-thread-myosin machinery and dissipation within the thin raphe; ii) external interactions, which include all interactions outside the cell such as the temperature-dependent hydrodynamic drags, where temperature can alter the surrounding viscosities.

To quantify the effects of these regimes, we isolate internal contributions from external effects by introducing 0.5 *μ*m polystyrene tracer particles beneath the diatom (ventral region). Interestingly, these tracer particles move opposite to the diatom’s travel direction, indicating attachment to mucilage threads connected to the internal myosin machinery (Fig. 2f and Movie S5). This opposing movement contrasts with previous studies where tracer particles attached to dorsal regions showed no correlation with diatom movement^82^. The movement of ventral tracer particles mirrors the internal motion of myosin motors and maps the kinetics of the mucilage-thread-myosin machinery right outside the raphe. Quantifying particle speeds reveal that when particle movement is present, diatom movement slows down significantly (Fig. 2g), demonstrating that particles attached to the mucilage disrupt traction and inhibit forward motion. Additionally, we notice that some tracer particles aggregate into clusters of varying sizes. Their movement speeds show no correlation with cluster size (Fig. 2h). This invariant speed across different loading forces suggests that the diatom maintains a uniform myosin motility rate within a certain loading regime. Thus, we consider the movement speed of the tracer particles to be a reasonable approximation of the internal speed of the myosin-mucilage threads involved in diatom motion.

Comparing the average speeds of tracer particles with diatoms without particle attachment shows a significant velocity reduction (Fig. 2i). This velocity loss persists across the entire temperature range explored and intensifies as temperature decreases, reaching approximately 80% of velocity loss at temperatures below −5 °C (inset of Fig. 2i). This velocity reduction is likely due to external drags affecting the diatom more than the smaller particles. The temperature-dependent loss highlights the distinct impact of the external drags on diatom motility. Subsequently, we examine how a gliding diatom interacts with the environments.

Gliding motility necessitates close proximity to a substrate. Between the diatom surface and the substrate there is a gap filled with a thin film of mucilage solution (0.5 - 1 *μ*m^83^), which places the system within the lubrication regime and contributes substantially to hydrodynamic drag. This drag effect has been largely overlooked in past analyses^79,84^.

Our observations reveal that during smooth movement, ventral tracer particles migrate from the diatom’s tip toward the center, stall, and then move at a similar speed as the diatom, as shown in the kymograph in Fig. 3a. This pattern suggests that particles attached to the mucilage initially move with myosin motors along actin cables but later become passively dragged, indicating when the mucilage disconnects from the myosin. The disconnection near the cell’s center is likely due to a central discontinuity in the raphe (Fig. 3a). Although actin cables are continuous, allowing continuous movement of myosin (Fig. 2c), the raphe discontinuity interrupts movement of mucilage threads. The mucilage accumulates near the middle and eventually diffuses in the surrounding water. The discontinuous geometry suggests a division of the ventral region into a leading edge, where active ‘driving’ occurs via mucilage-thread-myosin machinery, and a trailing edge, where ‘dragging’ effects are predominantly observed.

**Figure 3.**
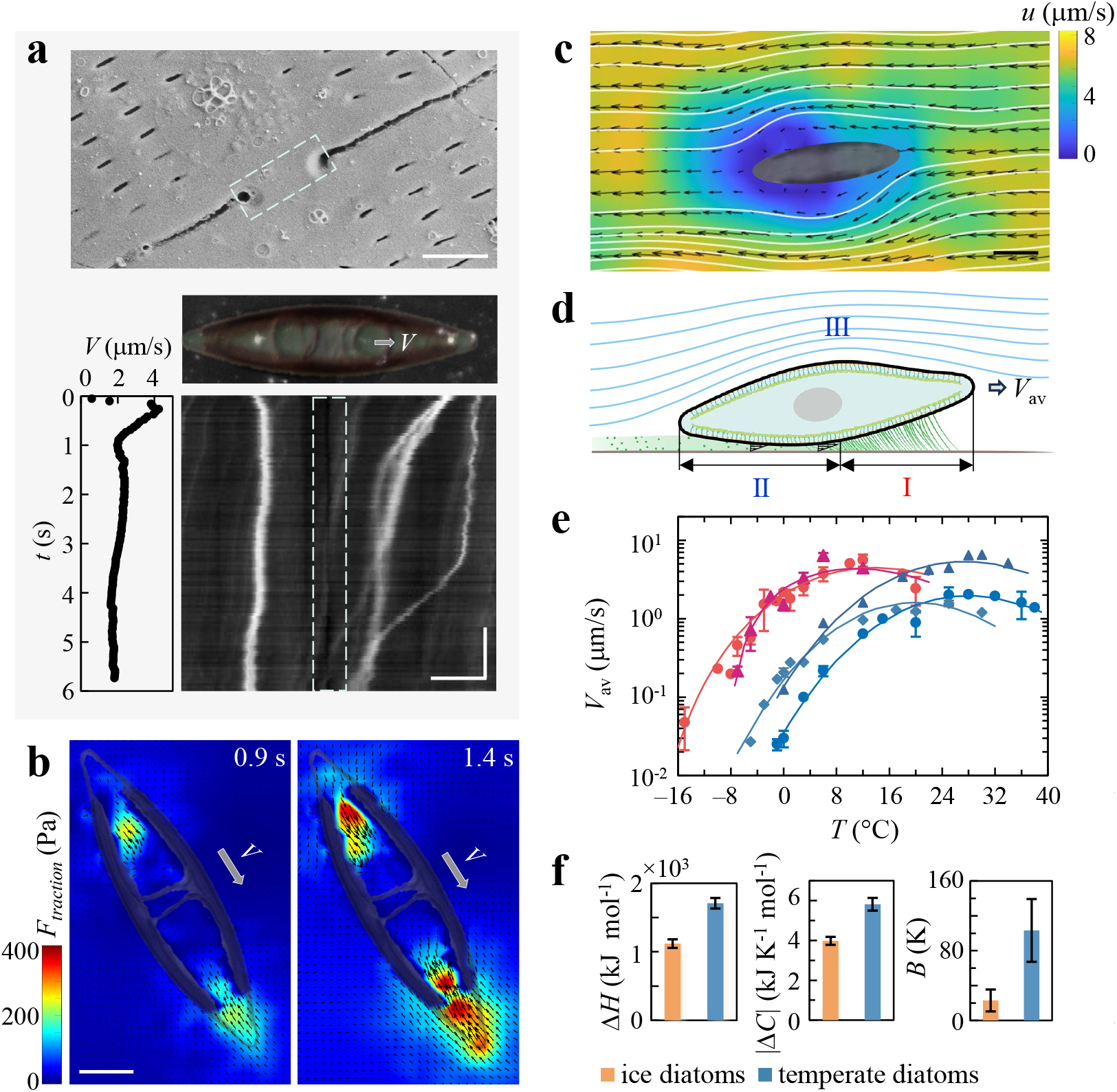
A thermo-hydrodynamic model incorporating internal and external factors illustrates ice diatoms’ strategies for enhanced gliding motility in cold environments. (**a**) Upper panel: SEM image of an ice diatom showing a raphe discontinuity (light blue boxes) dividing the ventral region into leading and trailing edges. Scale bar: 1 *μ*m. Lower right panel: Kymograph from motility videos showing tracer particles latched onto the ventral raphe; particles on leading and trailing edges are highlighted above (white). Lower left panel: Diatom’s velocity over time. In the diatom’s frame of reference, leading-edge particles moving backward along the raphe indicate active force generation. Particles near or in the trailing edge are passively dragged, demostrating that mucilage released into the surrounding water contributes to the lubrication film. Scale bars: 10 *μ*m (horizontal) and 1 s (vertical). (**b**) Two dimensional traction force microscope images taken over time reveal driving forces consistently applied in the leading edge and competing drag forces experienced in the trailing edge. Scale bar: 10 *μ*m. (**c**) Particle image velocity (plotted in diatom frame of reference) depicts ambient hydrodynamic drag experienced by a gliding ice diatom. Arrows indicate direction of flow and colors represent velocity magnitude. White lines depict streamlines. Scale bar: 20 *μ*m. (**d**) Schematic highlighting three regimes governing gliding motility: (I) Internal driving region from mucilage-thread-myosin machinery, and external drag regions from (II) ventral viscoelastic drag, and (III) ambient Stokes drag. (**e**) Comparative analysis of average gliding speeds for ice and temperate marine diatoms across varying temperatures, with lines representing the best fits to Eq. 1: 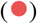 ice *Navicula sp*., 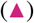 ice *Pluerosigma sp*., 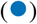 temperate *Navicula sp*., 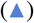 temperate *Pluerosigma sp*., 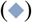 temperate *Pinnularia sp*., showing that the model captures the primary factors governing gliding speeds. (**f**) Comparison of activation enthalpy (Δ*H*), changes in heat capacity (Δ*C*) and sensitivity of mucilage viscosity to temperature (*B*) for both ice and temperate marine diatoms. The differences highlight the adaptive strategies allowing ice diatoms to glide in extreme cold.

Traction force microscopy (TFM) further elucidates these effects. Analysis of sustained movement of cells shows that traction forces opposing the diatom’s movement (attributable to the mucilage-thread-myosin machinery) are predominately localized at the leading edge, as shown in Fig. 3b and Movie S6. These forces facilitate the diatom’s forward motion. Conversely, traction forces aligned with the movement direction are primarily at the trailing edge, creating a ‘tug-of-war’ effect. Rather than solely due to bidirectional myosin activity^79^, this phenomenon is more likely due to viscoelastic stresses from passively dragged mucilage, as evidenced in Fig. 3a where tracer particles at the trailing edge move synchronously with the diatom.

The magnitude of ventral viscoelastic drag depends on the mucilage rheological properties. Given that the timescale of mucilage diffusion is comparable to the characteristic time associated with the gliding speed (see Supplementary Information), the mucilage layer persists beneath the moving diatom. We consider the ventral mucilage solution as viscoelastic fluid and model it using a Maxwell viscoelastic model: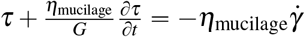, where *τ* is the shear stress, *G* is the elastic modulus, and *η*_mucilage_ is the zero-shear-rate viscosity of mucilage solution. Under steady-state conditions, the shear stress is proportional to an effective viscosity. Estimated viscoelastic drag, ranging from 200 to 2,000 pN at 1 *μ*m */* s, is on the same order of magnitude as the propulsive force generated by ice diatoms (1,200 pN) (estimated from TFM; see Supplementary Information). Therefore, ventral viscoelastic dissipation significantly affects diatom motility and contributes to reduced velocity during force transmission from internal motors to overall movement.

To complete the hydrodynamic picture, we further examine flow dynamics above a gliding diatom using particle image velocimetry (PIV) (Fig. 3c). The ambient fluid exhibits Stokes flow, exerting additional resistance on the diatom frustule.

Since these mechanisms are generalisable to both ice and temperate diatoms, we develop a model incorporating internal and external temperature-dependent terms to compare species and reveal cold-adapted strategies (Fig. 3d). Assuming steady-state motion, we balance the internal force (propulsion from myosin motors and friction from mucilage threads moving through the raphe) with external forces (ventral viscoelastic drag and Stokes drag): *F*_inner_ = *F*_ventral_ + *F*_stokes_, which yields an expression for the average velocity of a gliding diatom (details in Supplementary Information):

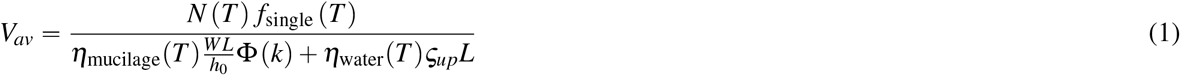

where *T* is the temperature, *N* (*T*) is the number of myosin motors activated and engaged during motility, *f*_single_ (*T*) is the force produced by single mucilage-thread-myosin machinery, *η*_mucilage_ (*T*) and *η*_water_ (*T*) are the temperature-dependent viscosities of mucilage and water, respectively, *L* is the length of diatom, *W* is the width of the diatom, and Φ (*k*) is a geometric factor dependent on the ratio of the front gap size to the back gap size: *k* = *h*_1_ *h*_0_.The gap is defined as the distance between the ventral side of the diatom and the substrate.

We first account for temperature-dependent internal factors: *N* (*T*) is governed by enzyme-catalyzed kinetics intrinsic to the myosin-actin interaction cycle, and *f*_single_ (*T*) is controlled by conformational changes in myosin molecules^86,87^. These biological processes depend on temperature. A fundamental theory that describes the temperature dependence of such biological processes is the Eyring-Evans-Polanyi (EEP) transition state theory^88,89^. Based on EEP and under the assumption that the temperature dependence arises mainly from entropy changes, we model the internal force as^90^: 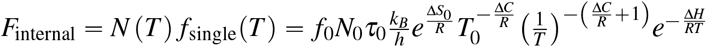, where *N*_0_, *f*_0_, *τ*_0_ and Δ*S*_0_ are the activated number of myosin motors, the force per motor, the characteristic time of the mechanochemical cycle, and the entropy change, respectively, all evaluated at the reference temperature *T*_0_ (typically 293K). *k*_*B*_, *h* and *R* are Boltzmann’s constant, Planck’s constant, and the gas constant. Δ*C* is the change in heat capacity. Δ*H* is the activation enthalpy.

We account for the impact of temperatures on environmental factors by incorporating the temperature-dependent viscosity of seawater, *η*_water_(*T*)^91, 92^, and modeling the mucilage viscosity using the Vogel-Fulcher-Tammann (VFT) equation^93^: 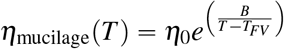, where *η*_0_ is the reference zero-shear-rate viscosity, *T*_*FV*_ is the critical glass transition temperature, and *B* characterizes the sensitivity of mucilage viscosity to temperature changes.

Based on temperature dependent gliding motility of both ice diatoms and temperate diatoms, we fit Eq. 1 to our data (see Supplementary Information; parameters listed in Table S3). The current model effectively describes the experimental data for both ice and temperate diatoms, as shown in Fig. 3e. The model captures the primary physical phenomena governing temperature-dependent motility.

The parameters from the EEP model (Δ*C* and Δ*H*) and the VFT model (*B*) provide physical intuition for predicting the strategies that ice diatoms might have evolved to enhance motility under freezing conditions. Compared to temperate species, ice diatoms exhibit a smaller change in heat capacity (Δ*C*), suggesting they are better adapted to maintaining internal stability against temperature fluctuations (Fig. 3f). This stability is advantageous in consistently cold environments where large internal energy fluctuations can be detrimental. Additionally, a lower activation enthalpy (Δ*H*) reduces the enthalpic barrier, enhancing energy efficiency at freezing temperatures to sustain internal driving dynamics.

In response to the external hydrodynamic effect in extreme conditions, ice diatoms show reduced sensitivity of mucilage viscosity to temperature variations, indicated by a smaller *B* (Fig. 3f). Assuming similar viscosities at room temperature, the mucilage of ice diatoms increases in viscosity more slowly as temperatures decrease compared to that of temperate diatoms. This adaptation allows ice diatoms to maintain higher mucilage fluidity and effectively reduce resistance forces, thereby facilitating enhanced motility in freezing conditions.

### Temperature-dependent motility in gliding population

The dynamics of ice diatoms’ gliding motility extends beyond simple considerations of average velocity, as shown in Fig. 4a. Our analysis of the statistical distribution of diatom motility reveals two distinct regions in the probability density function: a sharp decline at low speeds followed by a secondary peak and a subsequent decline at higher speeds (Fig. 4a). This distribution shows a remarkable dependence on temperature. At lower temperatures, the secondary peak is less pronounced, largely obscured by the prevalence of non-motile diatoms, resulting in a distribution that mostly decays. As temperatures increase, this secondary peak becomes more distinct, and the speeds at which this peak occurs also increase. Concurrently, as temperature rises, we observe a broadening range of speeds. This indicates that individual diatoms display more variability in their motility at increased temperatures.

**Figure 4.**
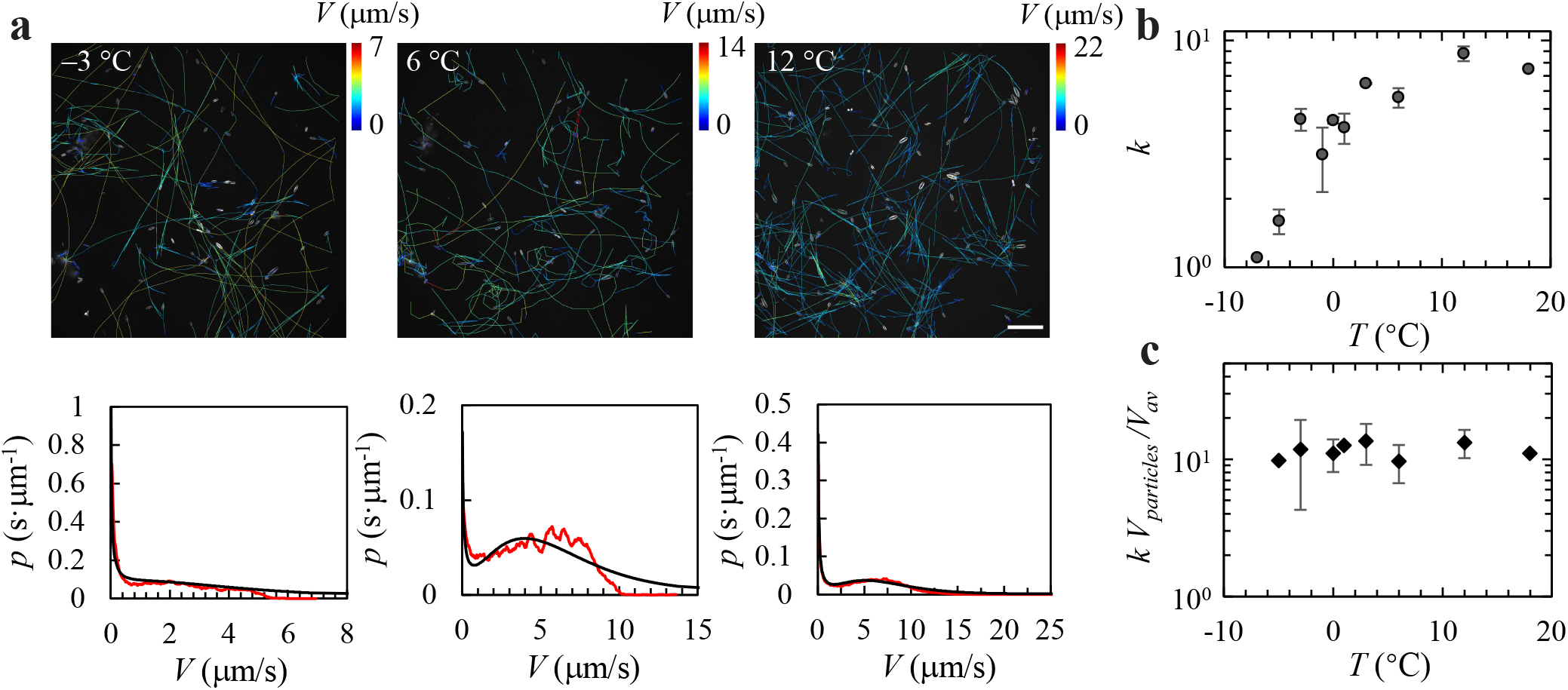
Temperature-dependent motility patterns in diatom populations. (**a**) Trajectories and gliding speed distributions on a glass surface at − 3 °C, 6 °C, and 18 °C, displayed from left to right. Upper panel: diatom trajectories. Scale bar: 200 *μ*m. Lower panel: the corresponding probability density functions, *p*, of gliding speeds *V* in red, fitted with a combination of Chi-squared and Gamma distributions in black. (b) Evolution of the shape parameter *k* from the Chi-squared model, illustrating a shift from skewed to more symmetric, bell-shaped distributions as temperature increases. (c) Normalization of *k* by the velocity ratio *V*_*av*_/*V*_*particle*_, showing how motility diversity correlates with efficiency across different temperatures.

These patterns deviate from a Gaussian distribution, which is typically indicative of random and uncorrelated movements. The absence of a Gaussian profile suggests that diatom motility is influenced by more complex internal and external factors rather than being purely stochastic. By analyzing the mean square displacement, we establish that their motion is super-diffusive, closely approximating ballistic movement (see Supplementary Information and Fig. S5). This implies that, once set in motion, ice diatoms maintain their trajectory over longer timescales than would be expected under normal diffusive conditions.

The statistical distribution of ice diatoms’ motility aligns well with a model combining Gamma and Chi-squared distributions, as shown in Fig. 4a:

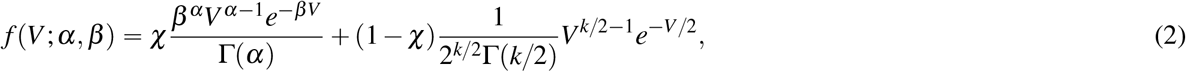

where *α* is the shape parameter, *β* is the scale parameter, Γ (*α*) and Γ (*k*/2) are the gamma functions, *k* is the degree of freedom, and *χ* is the weight coefficient. The second peak in the speed distribution, which corresponds to the most prevalent motility speeds among the active cells, is primarily described by the Chi-squared distribution (see Supplementary Information and Fig. S6). The Chi-squared distribution represents the squared velocity components of directional movement, consistent with the observed super-diffusive behavior. The spread of the motility distribution is characterized by the degrees of freedom, *k*. The value of *k* increases with temperature, reaching a plateau at around 12 °C, as shown in Fig. 4b. We present *k* only at temperatures down to 7 °C, because, below 7 °C, water freezes rapidly (within 1–3 minutes), leading to limited observation time and insufficient statistical information.

The increase in *k* indicates not just a simple speed variation, but a strategic diversification in movement patterns. Such diversification could serve as a bet-hedging mechanism against environmental unpredictability, allowing diatoms to exploit a wider range of microhabitats and resource patches.

Interestingly, the degree of freedom *k*, which characterizes the speed diversity, can be rescaled by the efficiency ratio of internal to external speeds, as shown in Fig. 4c. The rescaled result is a constant, which indicates that ice diatoms optimize their energy expenditure relative to the external conditions they encounter. As diatoms encounter less resistance, or as their internal mechanisms become more efficient, they can afford a greater diversity in their movement strategies with respect to energy expenditure. The interrelationship between temperature, motility diversity, and efficiency ratios reveals an adaptive strategy. It suggests that ice diatoms are not merely responding passively to temperature changes but are actively modulating their motility strategies in a way that maximizes their ecological success under varying thermal conditions.

## Discussion and Outlook

The gliding motility of ice diatoms in sea ice suggests finely tuned adaptations to their cold, dynamic habitat, highlighted in two aspects: distinct interaction with icy environments and resilience to cold temperatures.

The unique ability of ice diatoms to glide on ice, enabling them to thrive in conditions that immobilize other marine diatoms, correlates with their particular ability to adhere to ice through mucilage. The ice adhesion ability of ice diatoms, revealed and quantified by our shear flow assays, could be potentially enabled by well-known ice-binding proteins (IBPs) in mucilage threads. IBPs have been widely identified in ice diatoms^94^ to protect them from the cold by preventing recrystallization of ice^31–34^. However, the role of IBPs in aiding ice diatom mobility has not been documented. Certain IBPs exhibited in psychrophilic bacteria, like MpIBP_RIV, have demonstrated the ability to facilitate adhesion to ice^95^. Considering the hypothesis suggesting that IBPs in ice diatoms originate from psychrophilic bacteria^27,96,97^, it is plausible that ice diatoms might utilize similar mechanisms to adhere to ice. Further investigations of chemical components in the mucilage that contribute to the unique ‘skating’ ability of ice diatoms, potentially including IBPs, will deepen our understanding of the mechanism of adhesion in gliding motility on ice.

Additionally, the ice gliding motility suggests that the motor proteins of ice diatoms have adapted to low temperature conditions. A recent study has focused on identifying unique myosin motors in temperate diatoms^79^. More broadly, myosin motor-based single molecule studies have enabled us to understand the entire catalytic cycle of molecular motors walking^69^. These measurements have been mostly limited to room temperature studies and have not systematically explored colder regimes. We demonstrate the lowest temperature that myosin-driven motility can function. Future work will involve systematically dissecting the molecular machinery for these remarkable organisms that retain motility at sub-zero temperatures.

Ice diatoms have also evolved excellent adaptations to secure enhanced motility at cold temperatures by showing a systematic shift of their temperature responsive motility towards cold. This study integrates internal molecular machinery and external hydrodynamic factors to dissect mechanisms controlling the gliding speed of diatoms. The staining of mucilage trails and the visualization of the actin within the raphe region in ice diatoms confirm the internal basis of force generation in ice diatom motility^60,79^. The observations of particle movements in the ventral region of gliding diatoms allow us to quantify the impact of viscoelastic drags, distinct from internal driving mechanisms on diatom motility.

Employing traction force microscopy (TFM), we demonstrate the interactions between gliding ice diatoms and their substrates in real time. The driving force appearing at the diatom’s front moves synchronously with the cell (Movie S6), suggesting dynamic adjustments in force generation by myosin motors during gliding. The observed ‘tug-of-war’ phenomenon primarily arises from the competition between the internal driving force and the viscoelastic drag from mucilage beneath the diatom. This finding highlights the significant role of external hydrodynamic effects in modulating diatom motility. While our analysis assumes a steady-state scenario, future work will incorporate time-dependent processes to account for elastic contributions.

The hydrodynamic effects not only lead to a loss of output speed from the myosin motor but also tune the instantaneous dynamic behavior of diatoms in response to the environment. The observed tilt angle (Fig. 2c) reflects kinematic adjustments between the driving torque generated by the internal myosin motor and the counteracting hydrodynamic torques^78^. In studies of elliptical particles moving near walls, hydrodynamic torques can lead to tumbling behavior, characterized by a continuous change of the tilted angle^98^.

In conclusion, our study reveals a distinct aspect of how ice diatoms navigate sea ice, bridging ecology and cell physiology. Ice diatoms present ice gliding motility, which is absent in temperate counterparts. The unique ice gliding ability allows ice diatoms to search for optimal light, salinity and nutrient conditions within a dynamically changing landscape. The cold-adapted strategies predicted by our model provides insights for future research on how these psychrophilic microorganisms optimize motility under cold stress, including adaptations in motor protein conformations and rheological properties of mucilage. The observed temperature-dependent distributions in population motility reflect not just physical responses but strategic population adaptations. By linking movement statistics to biological energy expenditure and environmental interactions using a scaling analysis, we suggest that ice diatoms actively modulate energy utilization and movement strategies to optimize motility at the population level.

The discovery of ice gliding motility and the insights into energy-efficient movement can be integrated into ecological models to better predict phytoplankton dynamics within sea ice and inform adaptation strategies in other extremophilic microorganisms. These findings also add a dynamic perspective to climate prediction models for polar ecosystems under environmental change^99^.

## Methods

### Temperature-controlled fluorescent microscopy setup

Our studies utilize a facility-grade widefield fluorescent microscope (Squid, Cephla), custom-built with temperature-controlled moduli, as shown in Fig. 5a. This portable system is ideal for field studies due to its adaptability and mobility. Detailed configurations of the Squid microscope can be found in Ref.^100^.

**Figure 5.**
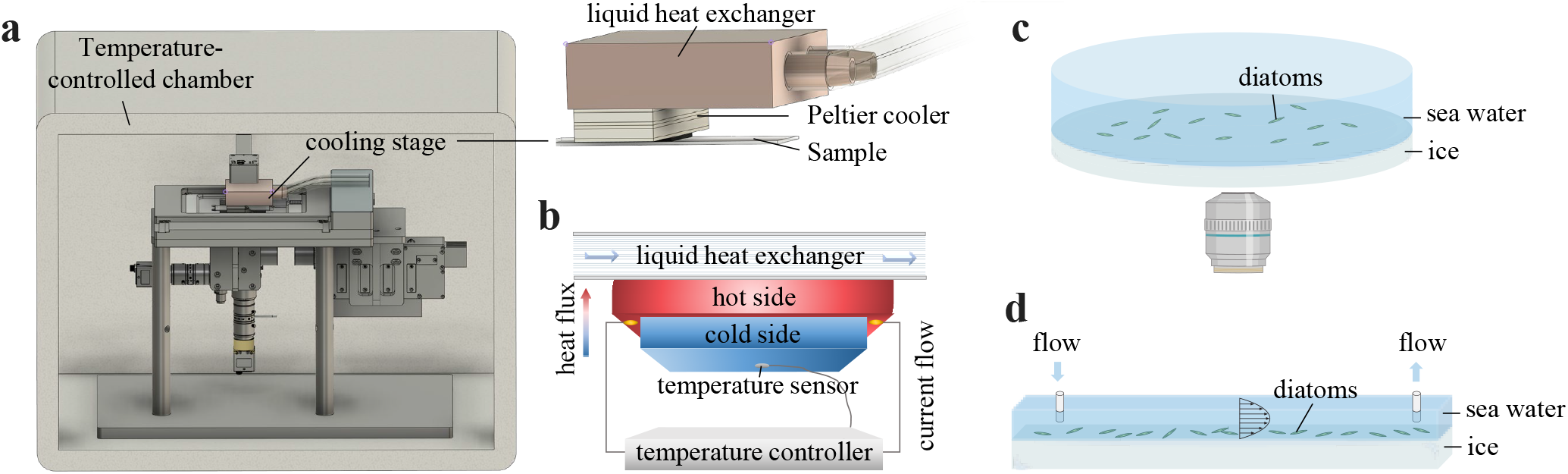
Experimental setups. (**a**) Sub-zero temperature-controlled fluorescent microscope equipped with a cooling stage and incubator. (**b**) Temperature feedback control of the cooling stage. (**c**) Setup for observing behavior of ice diatoms on ice surface. (**d**) Setup for quantifying the adhesion strength of ice diatoms on ice.

Temperature control in our experimental setup is achieved through two primary methods, as shown in Fig. 5a. The first method involves enclosing the entire microscope apparatus within a temperature-controlled chamber, providing stable conditions suitable for transmitted light imaging. The second method utilizes a cooling stage directly mounted on the microscope, designed for rapid temperature changes critical for studying temperature-responsive behaviors. This cooling stage enables swift temperature reductions from 20 °C to − 20 °C within a minute, facilitating precise control during fluorescent imaging. In this configuration, diatoms are observed via their autofluorescence at excitation wavelengths of approximately 470 nm and 640 nm. This dual approach allows imaging under either transmitted light or fluorescence, enhancing the flexibility and efficiency of our experiments.

Both the temperature-controlled chamber and the cooling stage are equipped with Peltier devices. Peltier devices operate by passing an electric current through a series of semiconductor elements, exploiting the Peltier effect to absorb heat from one side (the cold side) and release it on the opposite side (the hot side), as shown in Fig. 5b. The Peltier device used for the chamber is commercially assembled with a fan for heat dissipation (Laird: SAA-170-24-22). The Peltier device employed for the cooling stage (Same Sky: CP39255074H-2) is customized to enhance its cooling efficiency. We attached a liquid heat exchanger to the hot side of this Peltier device, allowing for more effective heat removal compared to air cooling alone (Fig. 5b). This customized configuration ensures sustained super-cooling capacity, enabling consistent maintenance of low temperatures during experiments.

To achieve precise thermal regulation and maintain stable temperature conditions, we utilize a feedback control system (Fig. 5b). A temperature sensor (Amphenol Thermometrics: SC30F103A) is affixed to the cold side to continuously monitor temperatures, with data relayed to a temperature controller (8TCM-X107). This controller compares the actual temperature against the desired setpoint and employs a PID (Proportional, Integral, Derivative) algorithm to finely adjust the cooling outputs, minimizing temperature deviations and optimizing environmental stability.

For experiments investigating temperature-responsive motility, samples are placed in an incubation chamber sandwiched between a glass slide and a cover slide. The cover slide is directly affixed to the cold side of the cooling stage via a thermal pad, which has a high thermal conductivity of 14.8 W/m·K, to facilitate rapid temperature transfer.

The maximum temperature differential between the applied temperature and the actual temperature experienced by the sample is within − 1 °C. Because the sample affixed to the cold side of the cooling stage does not allow transmitted light imaging, we record the motility of ice diatoms using their auto-fluorescence excited at 470 nm, capturing images at 0.05 - 0.5 frames per second. The average speeds of gliding motility are calculated from observations of 20 - 100 moving cells, with error bars representing the variability from independently repeated experiments.

### Systems designed for investigating ice diatom behaviors within ice environments

To investigate the behavior of ice diatoms both on and within sea ice, we design specific model systems to preserve the natural conditions as closely as possible. For observations of diatom activity on the ice surface, we initiate the experiment by freezing a layer of fresh water at the bottom of a petri dish to form an ice layer approximately 3 mm thick. Subsequently, sea water maintained at 0 °C containing suspended diatoms is gently poured into the petri dish. The entire assembly is then placed immediately into a 0 °C incubator. After a 30-minute period, diatoms settle at the interface between the ice and sea water (Fig. 5c), enabling monitoring of their behaviors on the ice surface under the microscope. Additionally, to observe diatom activity within the ice, we develop ice microfluidic channels. These channels are fabricated to support studies of diatom motility embedded within the ice structure. Detailed methods and observations are provided in the Supplementary Information.

To evaluate the adhesion strength of diatoms on different substrates, we utilize flow assays within fluidic channels. Initially, a solution containing suspended diatoms is injected into the channel. After diatoms settle on the substrate, sea water is pumped through the channel at progressively increasing flow rates. The flow rates are controlled using a peristaltic pump (Kamoer DIPump). We obtain the critical flow rate *Q*_*c*_ needed to detach diatoms from the surface, and use the corresponding critical shear rate 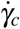 to characterize the adhesion strength. 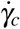 is taken as the maximum shear rate on the bottom wall of a rectangular duct in a laminar flow: 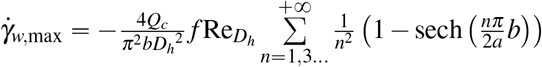, where *a* and *b* are the height and width of channels, respectively, *L* is the channel length, 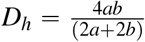 is the hydraulic diameter, 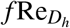 is Poiseuille number^101,102^.

For experiments of measuring adhesion strength on glass surfaces, the fluidic channels we use are configured with dimensions of 0.8 mm depth and 5 mm width (ibidi, *μ*-Slide). Conversely, assessing adhesion strength on ice requires the development of specialized ice channels. We fabricate channels with an initial depth of 8 mm and a width of 4 mm. Fresh water is injected to fill only the bottom layer of these channels, which is then frozen to form a 4 mm thick ice layer. The resulting channel’s depth is reduced to approximately 4 mm, with the ice forming the bottom surface where diatoms settle, as depicted in Fig. 5d. These channels, measuring 60 mm in length, are designed to allow the full development of the laminar flow. An observation window is strategically placed at the midpoint of the channel’s length and width to minimize boundary effects.

### Arctic ice cores post-processing in the field

At each Arctic ice station, ice cores are segmented into sections measuring 13 cm in length and 10 cm in diameter. We collect segments containing the biota layers from each ice core and place each of them into a large volume of 1 *μ*m-filtered seawater at an approximately 1:5 dilution. This mixture is maintained at 3 °C until it completely melts, allowing the diatoms to become fully suspended. Once they settle at the bottom of the container, we collect the sediments. These sediments are then used for further experiments, which include our direct observations of ice interactions in the field or culturing in the laboratory.

### Culture of ice diatoms and temperate diatoms

Using a dissection microscope, we isolate wild species of ice diatoms, specifically *Navicula sp*. (isolated from Ice Station 87 (71.67° N, 164.02° W) in the Chukchi sea). Initially, approximately 100 cells are isolated and washed six times to ensure maximal cleanliness. The isolated cells are then cultured in filtered seawater using F/2 medium under controlled conditions: 2 °C with a 12-hour light/12-hour dark cycle.

For comparative experiments, cultures of temperate diatoms including *Navicula sp*. (UTEX B SP11), *Pinnularia sp*. (UTEX B 679), *Navicula arenaria var. rostellata* (DCG 1006), and *Pleurosigma sp*. (DCG 1052) are obtained from the Culture Collection of Algae at the University of Texas at Austin and the Belgian Coordinated Collections of Microorganisms. These diatoms are grown in the same medium as previously described. The cultivation conditions for *Navicula sp*. and *Pinnularia sp*. are maintained at 20 °C, while *Navicula arenaria var. rostellata* and *Pleurosigma sp*. are cultivated at 14 °C, all under a 12-hour light/12-hour dark cycle.

### Sample preparation for fluorescent and SEM imaging

To investigate mucilage extrusion by ice diatoms, we place 1 ml of an ice diatom suspension in a glass dish, allowing the diatoms to glide on the substrate for 12 hours. We then add Wheat Germ Agglutinin (WGA) conjugated with Alexa Fluor 555 (Thermo Fisher Scientific) to the solution at a 1:200 dilution. The samples are incubated at 2 °C for 48 hours before imaging.

We further detect whether actin cytoskeletons exist in ice diatoms. The ice diatoms are fixed in 4% paraformaldehyde (PFA) for 1 hour at room temperature and washed three times with a 1:1 mixture of 1x PBS and filtered seawater. The final wash is performed using only 1x Phosphate Buffered Saline (PBS). Phalloidin conjugated to Alexa Fluor 546 (Thermo Fisher Scientific) is added at a 1:200 dilution and incubated for 1 hour at room temperature. Excess stain is removed by washing with 1x PBS, and samples are mounted in 80% glycerol for imaging.

Fluorescent staining is visualized using an inverted laser scanning confocal microscope (LSM 780/FLIM, Zeiss) equipped with a PLAN APO 63x/1.46 Oil objective lens (Zeiss). We excite WGA Alexa Fluor 555 and phalloidin Alexa Fluor 546 at 561 nm, detecting emissions with a 566/594 nm bandpass filter. Image processing is conducted using both Zen software and Fiji (ImageJ).

For SEM imaging, ice diatoms are fixed in 2.5% glutaraldehyde. After fixation, the samples are washed three times with 0.1 M sodium cacodylate buffer (pH 7.4) to remove excess glutaraldehyde. Subsequently, the samples undergo post-fixation in 1% osmium tetroxide (OsO_4_) for 1 hour under a fume hood. A graded series of ethanol solutions (50%, 70%, 95%, and 100%) are used for dehydration, with an extended incubation at 70% ethanol for 1 hour to enhance sample integrity. Finally, the samples are dried using hexamethyldisilazane (HMDS), a critical point drying technique that preserves the fine cellular structures by preventing surface tension effects during air drying.

### Traction force microscopy of ice diatoms on elastic polyacrylamide (PAA) substrates

Polyacrylamide (PAA) gel substrates are prepared to measure the traction forces exerted by ice diatoms. A prepolymer solution is formulated by mixing 5% (wt/vol) acrylamide and bis-acrylamide at a ratio of 19:1, which yields a gel with an approximate Young’s modulus suitable for cellular traction measurements^103^. To visualize substrate deformations, 200 nm fluorescent carboxylate-modified polystyrene beads (invitrogen, excitation 580 nm/emission 605 nm) are added to the prepolymer solution at a 1:200 dilution. Polymerization is initiated by adding 0.08% (wt/vol) ammonium persulfate (APS) and 0.1% (vol/vol) N,N,N’,N’-tetramethylethylenediamine (TEMED, Sigma-Aldrich) to the solution in a humidified chamber to prevent evaporation.

Immediately after adding the polymerization initiators, 30 *μ*l of the prepolymer solution is pipetted onto a clean glass coverslip (No. 1.5). A second coverslip is gently placed on top to spread the solution into a thin, uniform layer. The gel is allowed to polymerize undisturbed for 30 minutes at room temperature. After polymerization, the gel sandwich is carefully transferred into 1x PBS, and one coverslip is gently removed to expose the gel surface without disrupting the embedded beads.

Ice diatom samples are introduced onto the prepared PAA gel substrate by gently pipetting a diatom suspension onto the gel surface. The samples are incubated for 15 minutes to allow the diatoms to adhere to the substrate. Imaging is performed using our customized temperature-controlled fluorescence microscope equipped with a 40x objective lens (NA = 0.75). Temperatures are maintained at 3 °C to replicate cold environmental conditions. Fluorescence images of the embedded beads and the diatoms are acquired at a frame rate of 15 frames per second.

Image sequences are processed using MATLAB. Bead displacement fields are calculated using the Digital Particle Image Velocimetry tools in MATLAB (PIVlab)^104,105^. Two-dimensional traction forces are reconstructed from the displacement fields using the Fourier Transform Traction Cytometry (FTTC) method as described by Sabass *et al*.^106^. A regularization parameter of 2.5 · 10^5^ is used to balance the sensitivity and noise amplification in the calculations. This value is chosen based on the spatial resolution of our displacement field and the frequency content of the data, with a maximum squared wave number *K*_*max*_^2^ 4 10^14^ m^−2^. It effectively reduces noise without oversmoothing the traction maps, ensuring accurate representation of traction forces exerted by the cells. The Young’s modulus (*E* ≈ 7, 800 Pa) and Poisson’s ratio (0.5 for incompressible materials) of the gel are incorporated into the calculations.

### Analysis of hydrodynamic effects and motility of diatoms

To examine the flow fields induced by the motion of diatoms, we add fluorescent carboxylate-modified polystyrene beads of 0.5 *μ*m diameter (invitrogen, excitation 505 nm/emission 515 nm) in the solution containing diatoms. These beads are dispersed in the solution at a concentration of 0.05 wt%. These tracer particle experiments are conducted at various temperatures down to − 6 °C. Below this temperature, the water freezes immediately because the tracer particles act as nucleation sites for ice formation. Observations are conducted using the customized temperature-controlled fluorescence microscope, employing a 40x magnification objective (NA = 0.75).

For imaging the flow field in the gap thickness direction, we focus on two distinct layers: the bottom layer, located just beneath the diatoms, and the top layer, positioned above the diatoms’ midline. Both the lower and upper flow fields are recorded at a frame rate of 2 - 5 frames per second. The two-dimensional velocity field for the upper flow layers is analyzed using PIVlab^104,105^.

Additionally, the motion of particles beneath the diatoms and the motility of the diatoms themselves are analyzed using the machine learning-based image processing tool ilastik for segmentation and TrackMate for tracking trajectories^107,108^.

## Data availability

All data needed to evaluate the conclusions in the paper are present in the paper, and/or the Supplementary Information.

## Acknowledgements

We thank the captain and crew of the R/V Sikuliaq and the entire science team for support. We thank R. Konte for help on the schematic of diatom gliding mechanisms. We thank G. Zhong for assistance on the actin staining protocol, and I. Ho and X. Mao for helpful discussions on validating the theoretical model. We thank the Cell Sciences Imaging Facility at Stanford University and the assistance provided by D. Lenzi and R. Yamawaki. Q.Z. K.R.A. and M.P. acknowledge support from National Science Foundation (NSF) (grant no. 135316). H.T.L. acknowledges support from the Stanford VPGE DARE fellowship and the NSF Graduate Research Fellowship Program (grant no. DGE-1656518). M.P acknowledges further support from HFSP, Moore foundation, NSF CCC (DBI-1548297), Schmidt Foundation, and Dalio Foundation.

## Author contributions

Q.Z. and M.P. designed the research. Q.Z., H.L. and M.P. developed field-customized temperature-controlled microscope. Q.Z. and K.R.A. collected samples in the Arctic. Q.Z. performed the experiments. Q.Z. and M.P. performed theoretical analyses. Q.Z. and M.P. analysed the data. Q.Z., H.T.L., and M. P. wrote the paper.

## Additional information

**Supplementary information** is available for this paper.

