## Supplementary Information for "Ice gliding diatoms establish record-low temperature limit for motility in a eukaryotic cell"

**The PDF file includes:**

Supplementary Information

Figs. S1 to S6

Tables. S1 to S4

Legends for Movies S1 to S6

**Other Supplementary Information for this manuscript includes the following:**

Movies S1 to S6

### Biodiversity of ice diatoms in the Arctic ice cores

During our Arctic expedition (TOTS2023) in the Chukchi Sea, we collected ice cores from 12 ice stations, see Table S1. Ice cores with high biomass density are observed at stations 15, 32, 61, 87, and 92. Diatoms extracted from these high biomass ice cores are suspended in water and examined under high magnification (10x, 20x and 40x) to assess their biodiversity. Based on morphological characteristics, we identify the genera and observe the diatom community is predominantly composed of pennate diatoms, including *Navicula sp.*, *Pleurosigma sp.*, *Nitzschia frigida*, *Entomoneis sp.*, *Pseudo-nitzschia* and *Fragilariopsis sp*, as shown in Fig. S1(a–f).

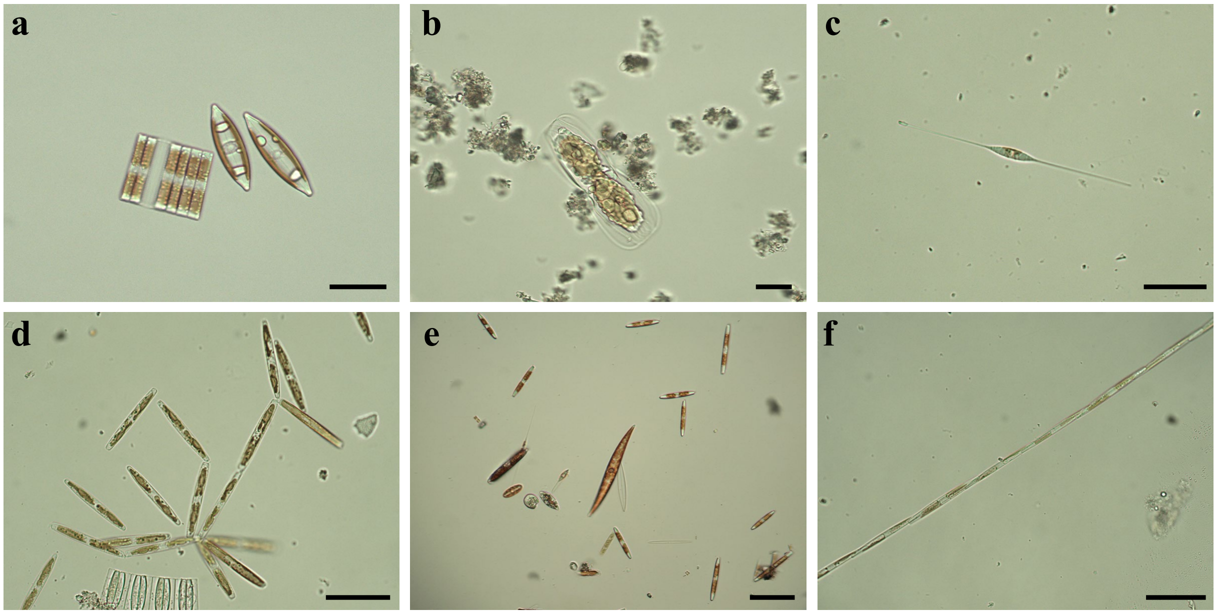


**Fig. S1. Genera of ice diatoms found in the Arctic ice cores.** The predominant diatoms are pennate-shaped, including (**a**) *Navicula sp.* and *Fragilariopsis sp.* (**b**) *Entomoneis sp.* (**c**) *cylindrotheca closterium* (**d**) *Nitzschia frigida* (**e**) *Pleurosigma sp.* and *Nitzchia sp.* (**f**) *Pseudo-nitzschia*. The scale bar is 50 $\mu$m.

### Investigation of ice diatoms' behavior in ice micro-channels

Sea ice is a porous medium filled with brine channels, which serve as habitats for ice diatoms[1–3]. To closely mimic this natural porous environment and investigate the activity of ice diatoms within ice, we develop an approach to fabricate ice micro-channels in the laboratory.

To create these micro-channels, we embed metal threads (diameter 100 $\mu$m) in water within a quasi-two-dimensional container, as shown in Fig. S2(a). After freezing the entire container at $-20$ °C, we remove the metal threads from the ice, resulting in the formation of ice micro-channels, as shown in Fig. S2(b). We then introduce the ice diatom *Navicula sp.* into the micro-channels and observe their activity under a microscope maintained at 0 °C with a magnification of 4x. The ice diatoms are able to glide freely within these ice channels, exhibiting behavior similar to that observed on two-dimensional ice surfaces. This setup allowed us to study their motility within confined ice environments, analogous to natural brine channels. The trajectory of an individual ice diatom is shown in Fig. S2(c) and Movie S2.

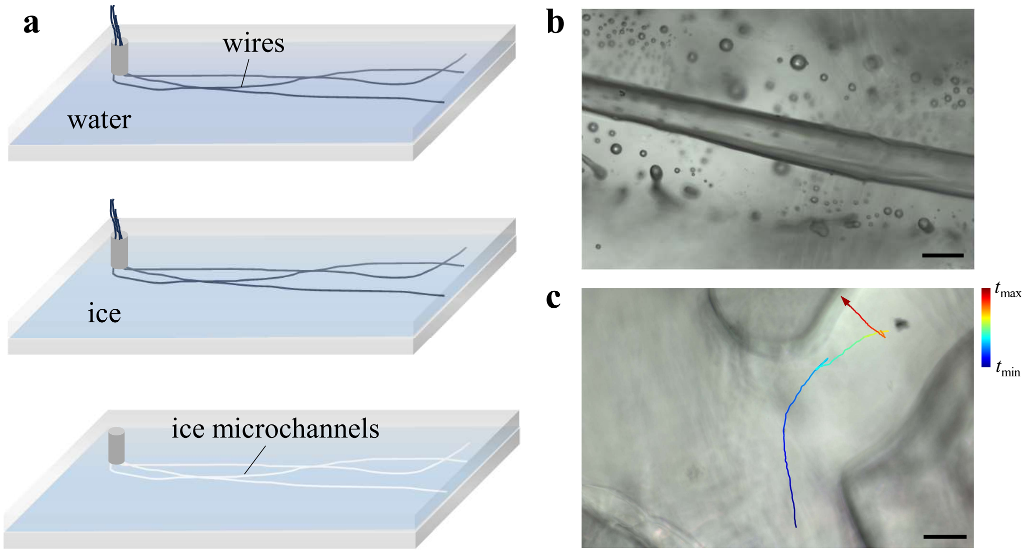


**Fig. S2. Ice microfluidic channels.** (**a**) Fabrication of ice micro-channels. Step 1 involves patterning using metal threads (top panel). Step 2 is the freezing process (middle panel). Step 3 is demolding (lower panel). (**b**) Example of an artificial micro-channel in ice. The scale bar is 200 $\mu$m. (**c**) Trajectories of ice diatoms moving within an ice micro-channel. The scale bar is 300 $\mu$m.

### Shift of temperature responsive motility for ice diatoms towards lower temperatures

To investigate the motility adaptations of ice diatoms to sub-freezing temperatures, we compare the temperature-dependent motility between ice diatoms and temperate marine diatoms. All diatoms exhibit similar temperature-dependent trends: as temperature increases, their average moving speed, $V_{\text{av}}$, normalized by the maximum velocity, $V_{\text{max}}$, increases, reaches a plateau, and then sharply declines to near zero at higher temperatures, as shown in Fig. S3(a). However, the temperature response curves for ice diatoms show a systematic shift toward lower temperatures, indicating striking cold adaptations in their motility (Fig. S3(a)).

To quantify this shift, we define characteristic temperatures for each curve. The minimum and maximum temperatures, $T_{\text{min}}$ and $T_{\text{max}}$, correspond to the temperatures below and above the points when the normalized velocity $V_{\text{av}}/V_{\text{max}}$ falls below 0.1. The optimal temperature, $T_{\text{optimal}}$, is the temperature at which the velocity reaches its maximum. Using interpolation where necessary, we obtain the values of $T_{\text{min}}$, $T_{\text{optimal}}$, and $T_{\text{max}}$ for both ice and temperate diatoms, as presented in Table S2.

Interestingly, the temperature response curves for different diatom species are collapsed onto a master curve by normalizing the deviation of temperature from the optimal temperature, $(T - T_{\text{optimal}})$, by the temperature range, $(T_{\text{max}} - T_{\text{min}})$. This rescaling reveals that both temperate and ice diatoms share a similar response to temperature with respect to motility, as shown in Fig. S3(b). However, ice diatoms have evolved a systematic shift toward colder temperatures, highlighting their specialized adaptations to freezing environments.

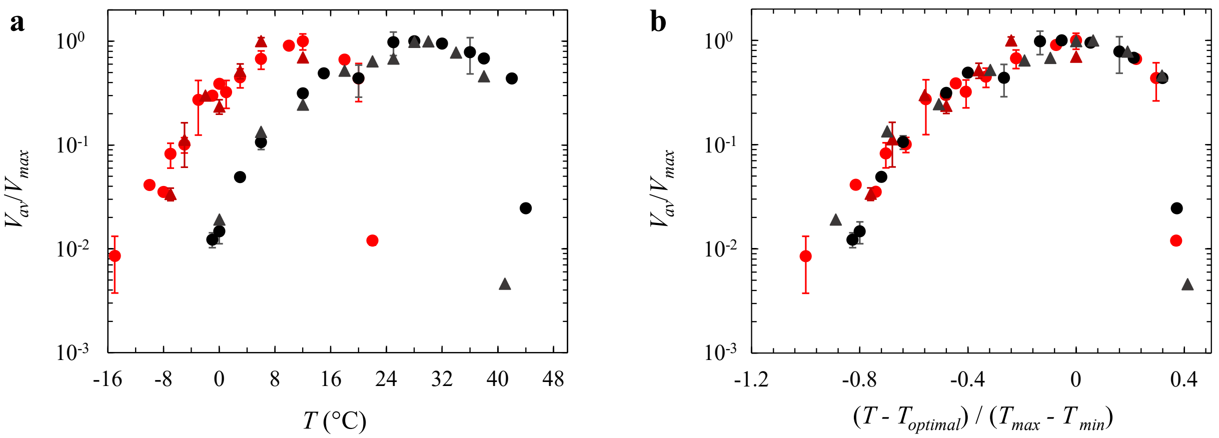


**Fig. S3. Comparison of temperature responsive motility between ice and temperate marine diatoms.** (**a**) Average moving speed of ice and temperate diatoms, $V_{\rm av}$, as a function of temperature, $T$. (**b**) Scaled master curve of $V_{\rm av}/V_{\rm max}$ versus $(T_{\rm av} - T_{\rm optimal})/(T_{\rm max} - T_{\rm min})$. (**a** - **b**) (●) ice *Navicula sp.*, (▲) ice *Pluerosigma sp.*, (●) temperate marine *Navicula sp.*, (▲) temperate marine *Pluerosigma sp.*.

### Temperature dependence of cellular motility and transport across the tree of life

To comprehensively assess the impact of temperature on motility and transport processes at cellular and molecular scales, we conduct an extensive literature review. We compile representative temperature ranges over which motility or transport is observed in various eukaryotic cells, prokaryotic cells, and motor proteins[4–28]. These findings are summarized in Fig. S4.

Our review reveals that cellular motility is generally confined to specific temperature ranges that reflect the organisms' environmental adaptations. Notably, as discussed in the main text (Fig. 1f), ice diatoms exhibit gliding motility at temperatures as low as $-15$ °C, representing the lowest recorded temperature limits for motility in eukaryotic cells. This exceptional adaptation highlights the unique mechanisms ice diatoms employ to maintain cellular functions under extreme cold conditions and underscores the importance of studying their motility strategies to understand life in polar environments.

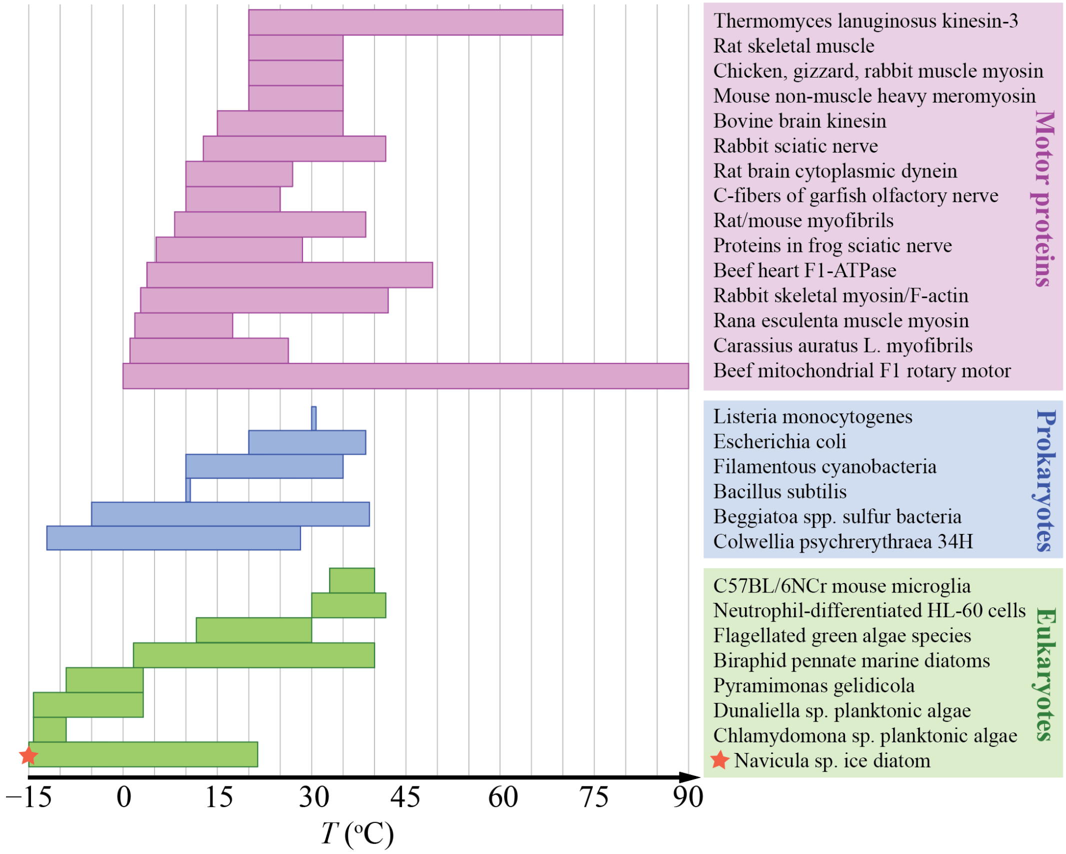


**Fig. S4. Recorded temperature dependence of cellular motility and transport.** Temperature ranges over which motility or transport processes have been recorded in eukaryotic cells, prokaryotic cells, and motor proteins[4–28]. This compilation highlights the impact of temperature on cellular and molecular motility across the tree of life, emphasizing that ice diatoms exhibit motility at the lowest recorded temperatures for eukaryotic cells.

### Modeling the temperature-dependent speed of diatoms

**Derivation of the theory.** The gliding speed of diatoms is governed by both internal machinery interactions (mucilage-thread-myosin machinery) and external hydrodynamic effects (ventral viscoelastic drag and ambient Stokes drag forces).

Internally, the motion originates from a combination of forces: the driving force exerted by myosin motors and the resistance resulting from the movement of mucilage threads through the raphe, collectively referred to as 'internal forces'. For each mucilage-thread-myosin component, the force is represented as $f_{single}(T)$, which combines the driving force from a single actomyosin motor and the friction of a connected mucilage thread moving in the diatom raphe. The friction between mucilage thread and raphe is considered as an interaction between two solid surfaces and assumed to be relatively insensitive to temperature within the physiological range. For the driving force from a single motor, although direct studies on diatom's myosins are limited[29], myosin motors across different organisms share conserved structural and mechanochemical properties[30, 31]. In muscle myosins, force generation has been shown to be an endothermic process and governed by the conformational change within the motor domain of the myosin head, with the force per actin-myosin interaction increasing with temperature[17, 32, 33]. Considering the conserved nature of myosin motor domains[30, 31], we hypothesize that force generated by diatom myosins exhibit similar temperature-dependent behaviors[25]. Therefore, $f_{\text{single}}(T)$ is temperature-dependent, and we express the impact of temperatures on $f_{\text{single}}(T)$ using Eyring-Evans-Polanyi (EEP) transition state theory[34, 35]:

$$f_{\text{single}}\left(T\right) = f_0 e^{\frac{\Delta S_f}{R}} e^{-\frac{\Delta H_f}{RT}}, \tag{S1}$$

where $f_0$ is force per myosin motor at the reference temperature $T_0$ (commonly 293 K). $\Delta S_N$ and $\Delta H_N$ are respectively entropy and enthalpy changes associated with force generation.

The number of activated myosin motors, $N(T)$, is also temperature-dependent[36] and can be derived from the general temperature-dependent model[37]:

$$N\left(T\right) = N_0 \tau_0 \frac{k_B}{h} e^{\frac{\Delta S_N}{R}} T_0^{-\frac{\Delta C}{R}} \left(\frac{1}{T}\right)^{-\left(\frac{\Delta C}{R}+1\right)} e^{-\frac{\Delta H_N}{RT}}, \tag{S2}$$

where $N_0$ is the activated number of myosin motors at the reference temperature $T_0$, $\tau_0$ is the characteristic time defining the mechanochemical cycle of myosin while activated at $T_0$, $k_B$ is the Boltzmann constant, $h$ is

Planck's constant, $R$ is the gas constant, $\Delta S_N$ is the change in entropy at $T_0$, $\Delta C$ is the change in heat capacity at constant pressure during activation, $\Delta H_N$ is the activation enthalpy, $T$ is the temperature.

The internal forces are proportional to $N$ and $f_{single}$. Combination of Eq. S1 and Eq. S2 gives:

$$F_{\text{internal}} = N\left(T\right) f_{\text{single}}(T) = f_0 N_0 \tau_0 \frac{k_B}{h} e^{\frac{\Delta S_0}{R}} T_0^{-\frac{\Delta C}{R}} \left(\frac{1}{T}\right)^{-\left(\frac{\Delta C}{R}+1\right)} e^{-\frac{\Delta H}{RT}}, \tag{S3}$$

where $\Delta S_0 = \Delta S_N + \Delta S_f$ and $\Delta H = \Delta H_N + \Delta H_f$.

It is interesting to note that the internal velocity of myosin motors, $V_{\text{internal}}(T)$, determined by the activity of myosin motors, is also temperature-dependent. It can be similarly derived from the general theory for temperature dependence in biology[37]:

$$V_{\text{internal}}(T) = l_0 \frac{k_B}{h} e^{\frac{\Delta S_0}{R}} T_0^{-\frac{\Delta C}{R}} \left(\frac{1}{T}\right)^{-\left(\frac{\Delta C}{R}+1\right)} e^{-\frac{\Delta H}{RT}}, \tag{S4}$$

where $l_0$ is a characteristic length from the deforming length of a myosin protein at reference temperature $T_0$.

Externally, the diatom experiences hydrodynamic effects, which include ventral viscoelastic drag associated with released mucilage and Stokes drag acting on the diatom's body. In the ventral region, we simplify the modeling by assuming a steady-state condition, so that the drag force is proportional to an effective viscosity. Given the small gap between a gliding diatom and the substrate, we model the ventral region as a lubrication problem and further consider the cell as a tilted block moving over a substrate. The gap between the diatom and the substrate varies from a largest value $h_1$ at the front to a smallest value $h_0$ at the rear end of the driven region. We approximate the gap height $h(x)$ in the driven region as linearly decreasing from $h_1$ to $h_0$ over the length of the diatom, $L$, and the lubrication force is given by[38]:

$$F_{\text{ventral}} = \eta_{\text{beneath}}(T) \frac{WL}{h_0} \left(\frac{1}{k-1}\right) \left(4 \ln k - \left(\frac{6\left(k-1\right)}{k+1}\right)\right) V_{\text{av}}, \tag{S5}$$

where $k = h_1/h_0$ is the ratio of the front gap size to the back gap size, $V_{\text{av}}$ is the average gliding speed of the diatom, $W$ is the width of the diatom, and $\eta_{\text{beneath}}(T)$ is the temperature-dependent viscosity of the fluid beneath the diatom.

To calculate the viscous drag, it is essential to characterize the local viscosity $\eta_{\text{beneath}}(T)$. This viscosity depends on whether the mucilage secreted by the diatom remains beneath it or diffuses into the surrounding water.

Initially, the mucilage is secreted by the cell and forms a highly entangled network that supports the diatom's motion. Over time, the mucilage is released and gradually diffuses into the water. We compare the timescales of diffusion and gliding to determine whether the mucilage predominantly remains beneath the diatom during motion. The diffusion time $t_{\text{diffusion}}$ over a characteristic length $\xi$ is given by $t_{\text{diffusion}} = \frac{\xi^2}{D}$, where $D$ is the diffusion coefficient of the mucilage. The characteristic gliding time $t_{\text{glide}}$ over the same length $\xi$ is $t_{\text{glide}} = \frac{\xi}{V_{\text{av}}}$. There, the ratio of diffusion time to gliding time is $\frac{t_{\text{diffusion}}}{t_{\text{glide}}} = \frac{\xi V_{\text{av}}}{D}$, where $\xi = W/2$ is the characteristic length scale based on the diatom's width $W$. Given that the primary components of mucilage are polysaccharides[29, 39], the diffusion coefficient $D$ is on the order of $10^{-14}\ \mathrm{m^2/s}$[40, 41]. Assuming represented values for $\xi = 15\ \mu\mathrm{m}$ and $V_{\text{av}} = 1\ \mu\mathrm{m/s}$, we find that $\frac{t_{\text{diffusion}}}{t_{\text{glide}}} \propto 10^2$. The diffusion time is much larger to the gliding time, indicating that the mucilage largely remains beneath the diatom during motion. Therefore, the local viscosity beneath the diatom is mainly contributed by the mucilage, and we have $\eta_{\text{beneath}}(T) = \eta_{\text{mucilage}}(T)$. Due to the high concentration of mucilage beneath the diatom, its viscosity exhibits significant temperature dependence. This behavior can be described by the Vogel-Fulcher-Tammann (VFT) equation, which is commonly used to model the viscosity of supercooled liquids and highly concentrated polymer solutions[42]:

$$\eta_{\text{mucilage}}(T) = \eta_0 e^{\left(\frac{B}{T - T_{FV}}\right)}, \tag{S6}$$

where $\eta_0$ is a reference zero-shear-rate viscosity, $T_{FV}$ is the Vogel temperature corresponding to a glass transition in polymers, and $B$ is a constant characterizing the sensitivity of mucilage viscosity to temperature.

In the Stokes drag region, the diatom experiences resistance as it moves through water. Approximating the diatom as a prolate spheroid moving longitudinally, the Stokes drag force is given by[43]:

$$F_{\text{Stokes}} = 16\pi\eta_{\text{water}}(T)e^3\left[\left(1 + e^2\right)L' - 2e\right]^{-1}V_{\text{av}}, \tag{S7}$$

where $\eta_{\text{water}}(T)$ is the viscosity of water[44, 45], $e = \sqrt{1 - \frac{W^2}{L^2}}$ is the eccentricity of the spheroid, and $L' = \ln\left(\frac{1+e}{1-e}\right)$. Assuming steady-state motion, the forces acting on the diatom are balanced:

$$F_{\text{internal}} = F_{\text{ventral}} + F_{\text{Stokes}}, \tag{S8}$$

Substituting (S3), (S5) and (S7) into (S8) and solving for $V_{\text{av}}$, we obtain:

$$V_{\text{av}} = \frac{\upsilon\left(\frac{1}{T}\right)^{-\left(\frac{\Delta C}{R}+1\right)} e^{-\frac{\Delta H}{RT}}}{\eta_{\text{mucilage}}(T)\beta_1 + \eta_{\text{water}}(T)\beta_2}, \tag{S9}$$

where $\beta_1 = \frac{WL}{h_0}\Phi(k))$ with $\Phi(k) = \left(\frac{1}{k-1}\right)\left(4\ln k - \left(\frac{6(k-1)}{k+1}\right)\right)$, $\beta_2 = \varsigma_{up} L$, $\varsigma_{up} = 16\pi e^3\left[\left(1+e^2\right)L' - 2e\right]^{-1}$ is defined as the Stokes coefficient, and $\upsilon = f_0 N_0 \tau_0 \frac{k_B}{h} e^{\frac{\Delta S_0}{R}} T_0^{-\frac{\Delta C}{R}}$. This expression encapsulates the interplay between internal enzymatic activity and external hydrodynamic forces affecting diatom motility.

To elucidate the temperature-dependent impacts of these factors on velocity, we rearrange (S9) and take the natural logarithm:

$$\ln(V_{\text{av}}) = \ln\left(\frac{\upsilon}{\beta_2}\right) + \ln\left(\left(\frac{1}{T}\right)^{-\left(\frac{\Delta C}{R}+1\right)} e^{-\frac{\Delta H}{RT}}\right) - \ln\left(\frac{\beta_1}{\beta_2}\eta_{0,\text{mucilage}} e^{\left(\frac{B}{T-T_{FV}}\right)} + \eta_{\text{water}}(T)\right)). \tag{S10}$$

In (S10), the Arrhenius-like dependence of ln($V_{\text{av}}$) on $1/T$ is modified by the temperature-dependent viscosities, reflecting both internal enzymatic kinetics and external hydrodynamic effects.

To determine the parameters in our model, we first isolate the internal parameters: the change in heat capacity $\Delta C$ and the activation enthalpy $\Delta H$. This is achieved by fitting the measured temperature-dependent speeds of tracer particles attached to the mucilage threads, which reflect the activity of the myosin motors, using (S4). Specifically, for ice and temperate *Navicula sp.*, temperate *Pluerosigma sp.*, and temperate *Pinnularia sp.*, we fit the internal velocity $V_{\text{internal}}(T)$ data to extract values for $\Delta C$ and $\Delta H$. For ice *Pluerosigma sp.*, since we did not have access to cultures in the lab to conduct tracer particle experiments, we constrain $\Delta C$ and $\Delta H$ within the bounds determined from the ice *Navicula sp.* results, considering that the enzymatic mechanisms are conserved across species. With the internal parameters determined, we return to (S10) to fit the overall diatom velocity $V_{\text{av}}$. The remaining parameters to be fitted include $B$, $\frac{\beta_1}{\beta_2}\eta_0$, $T_{Fv}$, and $c_1$, where $c_1$ is a dimensionless coefficient: $c_1 = \ln\left(\frac{\upsilon}{\beta_2}\right) - \ln\left(\frac{\beta_1}{\beta_2}V_0\eta_{0,\text{mucilage}}\right) + \ln\left(\left(\frac{1}{T}\right)^{-\left(\frac{\Delta C}{R}+1\right)} e^{-\frac{\Delta H}{RT}}\right) = \ln\left(\upsilon\left(V_0\beta_1\eta_{0,\text{mucilage}}\right)^{-1}\left(\frac{1}{T_0}\right)^{-\left(\frac{\Delta C}{R}+1\right)} e^{-\frac{\Delta H}{RT_0}}\right)$. $V_0 = 1\ \mu\text{m}$ is a representative velocity characterizing the magnitude of moving speed of most pennate diatoms. By fitting the experimental data for each species using (S10), we extract the remaining parameters. This comprehensive fitting allows us to disentangle the contributions of internal enzymatic activity and external hydrodynamic effects on diatom motility across temperatures. The fitted results are summarized in Table S3.

**Validation of the thermo-hydrodynamic model.** In our thermo-hydrodynamic model, we consider the average forces acting on the diatom by segmenting the flow field, rather than solving the entire flow field explicitly. While this approach simplifies the analysis, a more rigorous solution for an object moving parallel to a wall can be obtained using asymptotic methods that match the inner (analogous to lubrication effects) and outer (analogous to viscous drag) flow fields[46–48].

For a diatom translating parallel to a plane wall, we approximate the total hydrodynamic force in the direction of motion using the solution for a sphere, given by[47]:

$$F_x = -6\pi\eta a V_{\text{av}} \left[ \frac{8}{15} \ln\left(\frac{1}{\varepsilon}\right) + A + O(\varepsilon) \right] \tag{S11}$$

where $F_x$ is the total force acting on the sphere in the $x$-direction, $\eta$ is the dynamic viscosity of the fluid, $a$ is the characteristic length, taken as the average of the diatom's length and width $a = (H + L)/2$, $H \approx W$ is the height of the diatom, $\varepsilon = h/a$ is the dimensionless gap parameter, with $h$ being the characteristic separation between the sphere and the wall, taken as the average gap height $h = (h_0 + h_1)/2$, $A \approx 0.953$ is a numerical constant specific to the geometry. $O(\varepsilon)$ represents higher-order terms negligible as $\varepsilon \to 0$.

In (S11), the term involving the logarithm (inner solution) arises from the lubrication effect, capturing the significant increase in viscous resistance due to the thin fluid layer between the sphere and the wall. The constant term $A$ represents the outer solution, corresponding to the Stokes drag modified by the presence of the wall at larger distances. The ratio of the lubrication force to the Stokes drag force provides insight into the relative significance of these two effects: $\frac{\text{Lubrication Force}}{\text{Stokes Drag}} = \frac{\frac{8}{15}\ln\left(\frac{1}{\varepsilon}\right)46}{A}$.

In our model, we consider the lubrication and Stokes drag forces independently. To assess the validity of this approach, we compare our calculated forces with those obtained from an asymptotic method. Given ice diatoms with $L \approx 70\ \mu\text{m}$ and $W \approx 18\ \mu\text{m}$, the ratio between the inner solution and the lubrication force is approximately 1. This close agreement suggests that our approximation for the lubrication effect is reasonably accurate within the context of the assumptions made. Similarly, we compare the Stokes drag force from our model with the outer solution from the asymptotic analysis. The ratio between our calculated Stokes drag force and the outer solution is approximately 1.5. Given that both methods are approximate (our method through the segmented approach and the asymptotic method by applying solutions derived for spheres to diatoms), the discrepancies are expected. Additionally, we note that the Stokes drag is much less important than the lubrication drag in the diatom gliding system, because the viscosity in the ventral region (predominantly determined by the viscosity of mucilage solutions) is much larger than that in the ambient region (predominantly determined by the viscosity of water). We here consider both terms to describe the whole dynamic picture.

These comparisons indicate that our approach of treating the lubrication and Stokes drag forces independently yields results that are in reasonable agreement with an established method. The close agreements in the lubrication force and the Stokes drag force suggest that our method captures the primary physical mechanisms governing diatom gliding without the need to solve the entire flow field. While our current model provides a reasonable approximation, future research could focus on refining the estimation by incorporating more accurate geometrical representations of the diatom such as considering a spacial distribution of viscosities and introducing computational fluid dynamics simulations.

### Estimations of forces generated and experienced by gliding ice diatoms

We estimated the forces generated by ice diatoms using traction force microscopy measurements. The maximum traction stress produced at the front ventral region near the raphe is approximately 400 Pa. Considering an effective contact area of $A_{\text{traction}} \approx 3\ \mu\text{m}^2$, as determined from experiments, the maximum generated force is calculated as: $F_{\text{driving,max}} = \sigma_{\text{max}} \times A_{\text{traction}} \approx 1200$ pN.

For the ventral viscoelastic drag experienced by the diatom, we assume a steady shear state and approximate the viscosity of the mucilage solution to be 10 - 100 Pa s at freezing temperatures. When a diatom moves at a velocity of $V = 1\ \mu\text{m/s}$ with a gap of approximately $h_0 \approx 0.5\ \mu\text{m}$ between its ventral surface and the substrate, the viscous drag force can be estimated using the lubrication approximation: $F_{\text{ventral}} \propto \eta_{\text{mucilage}} \frac{V}{h_0} A_{\text{drag}}$, where $A_{\text{drag}} \approx 10\ \mu\text{m}^2$ is the area over which the drag is applied (estimated from the back region of the diatom near the raphe). Thus, the estimated ventral viscous drag force ranges from 200 pN to 2000 pN.

These calculations indicate that the ventral drag can be of the same order of magnitude as the propulsive forces generated by the diatom. Therefore, ventral viscous dissipation must be considered when analyzing the motility of ice diatoms, especially under freezing conditions where the mucilage viscosity is high.

### Super diffusive gliding motion of ice diatoms

The mean square displacements (MSD) of ice diatoms scale with time as $MSD \propto t^{\alpha_{MSD}}$, where the exponent $\alpha_{MSD}$ is greater than 1 and less than 2, as shown in Fig. S5. This indicates that their displacement over time increases faster than in Brownian diffusion (where $\alpha_{MSD} = 1$) but slower than in ballistic motion (where $\alpha_{MSD} = 2$)[49]. Therefore, ice diatoms exhibit super-diffusive behavior in their gliding motion. Notably, as the temperature decreases, the value of $\alpha_{MSD}$ remains nearly constant until temperatures drop below $-6$ °C. Furthermore, this super-diffusive behavior is consistently observed in both open and confined environments, regardless whether the substrate is ice or glass.

The persistent super-diffusive motion across various temperatures, substrates, and species suggests that ice diatoms possess a highly adaptive motility mechanism. This mechanism likely enables them to effectively navigate their heterogeneous icy habitats, which are characterized by fluctuating temperatures and spatial constraints[1,2].

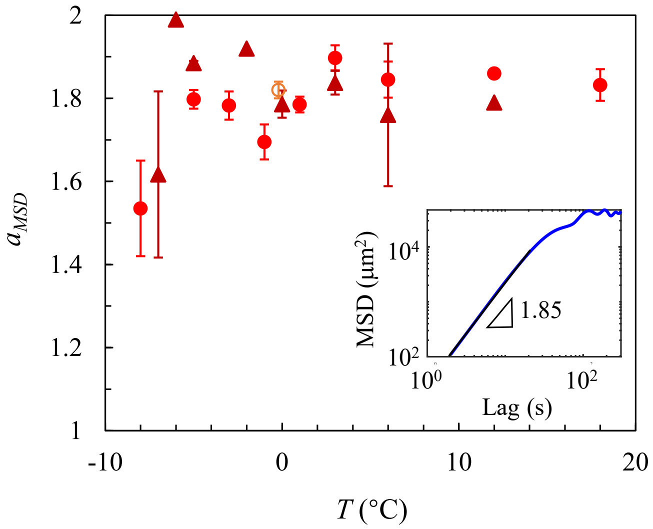


**Fig. S5. Mean square displacements of ice diatoms.** Ice diatoms exhibit super-diffusive behavior, characterized by mean square displacement slopes consistently greater than 1. This behavior is observed on glass substrates for (●) ice *Navicula sp.* and (▲) ice *Pluerosigma sp.*, and within ice micro-channels for (○) ice *Navicula sp.*. Inset: Mean square displacement showing the gliding motility of ice *Navicula sp.* at 3 °C.

### Statistics of ice diatom motility

Our analysis reveals that the motility patterns of ice diatoms across varying temperatures can be effectively modeled by a combination of Gamma and Chi-squared distributions (Fig. S6). The fitting parameters are shown in Table S4. This dual-model approach captures distinct aspects of their motility patterns across varying temperatures.

The Gamma distribution primarily describes the initial sharp decline at low speeds observed in the motility profiles, as shown in Fig. S6. The skewed nature of the Gamma distribution suggests a gradient in motility rates, reflecting variable energy expenditures and environmental resistances encountered by the diatoms. This indicates that some diatoms may navigate through their environments with varying efficiency, facing different degrees of friction, which contribute to the observed range of lower speed movements. Conversely, the second peak, which characterizes the most prevalent motility speeds among the active diatoms, is aptly modeled by the Chi-squared distribution (Fig. S6). This distribution is particularly suitable for representing the squared velocity components of directional movement, which aligns with the observed super-diffusive behavior of the diatoms (Fig. S5). As the temperature increases, this second peak becomes more pronounced and spreads out, indicating a broader range of higher velocity movements. The Chi-squared distribution effectively captures this dynamic, with the characteristic parameter $k$ increasing with temperature (as described in the main text).

The combination of these two statistical distributions provides a robust framework for understanding the complex dynamics of diatom motility, revealing how they adjust their locomotive behaviors in response to thermal variations in their habitats.

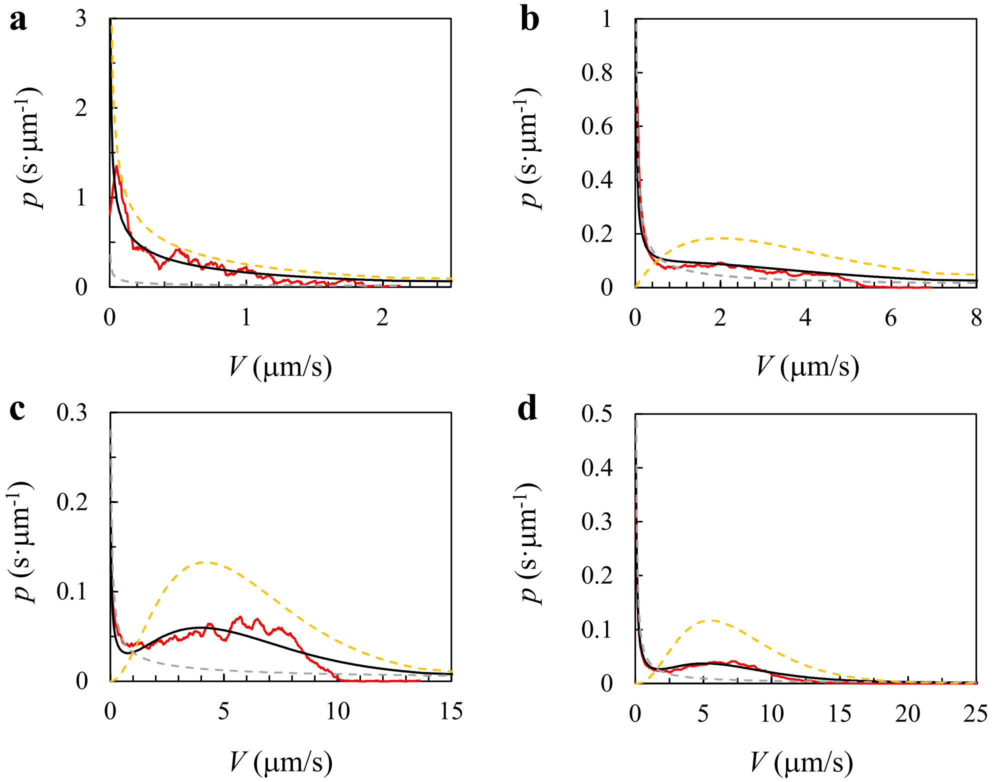


**Fig. S6. Fitting the statistics of ice diatom motility populations.** Contributions of Gamma (grey) and Chi-squared distributions (yellow) to fitting the probability density function in motility $p$ (black) at (**a**) $-7$ °C, (**b**) $-3$ °C, (**c**) 6 °C, and (**d**) 18 °C.

**Table S1.** Locations of ice stations and and chlorophyll information of ice cores of the 2023 Arctic Expedition.

| ice stations | latitude (°) | longitude (°) | highest Chl a ($\mu$g/L) | highest Chl depth (cm) | total thickness (cm) |
|---|---|---|---|---|---|
| 15 | 70.8893 | −161.571 | 174.4 | 124.0 | 124 |
| 21 | 71.2745 | −164.577 | 11.70 | 96.5 | 228.5 |
| 32 | 71.1787 | −163.023 | 119.28 | 85.0 | 85 |
| 38 | 71.4212 | −164.416 | 56.75 | 81.5 | 81.5 |
| 42 | 71.0638 | −165.000 | 8.57 | 45.0 | 102.5 |
| 56 | 71.2068 | −163.083 | 47.11 | 137.0 | 137 |
| 61 | 71.586 | −163.627 | 189.97 | 133.0 | 133 |
| 64 | 71.414 | −164.409 | 56.10 | 235.0 | 235 |
| 78 | 70.9157 | −163.110 | 19.11 | 75.0 | 82 |
| 83 | 71.3812 | −162.964 | 52.98 | 65.0 | 136 |
| 87 | 71.6722 | −164.020 | 125.49 | 118.0 | 125.5 |
| 92 | 71.1962 | −164.816 | 109.43 | 94.0 | 166 |

**Table S2.** Characteristic temperatures defining ice and temperate diatoms' temperature responsive motility.

| genus | ecotypes | $T_{min}$ (°C) | $T_{optimal}$ (°C) | $T_{max}$ (°C) |
|---|---|---|---|---|
| *Navicula sp.* | ice | $-7$ | 12 | 20 |
| | temperate | 5 | 30 | 42.5 |
| *Pleurosigma sp.* | ice | $-5$ | 12 | 20 |
| | temperate | 5 | 28 | 36.5 |

**Table S3.** Fitting parameters from the thermo-hydrodynamic model for temperature dependent gliding speed of ice and temperate diatoms.

| Genus | $\Delta C$ (J $K^{-1}$ $mol^{-1}$) | $\Delta H$ (J $mol^{-1}$) | $B$ (K) | $\frac{\beta_1}{\beta_2}\eta_0$ (Pa s) | $T_{Fv}$ (K) | $c_1$ |
|---|---|---|---|---|---|---|
| *Ice Navicula sp.* | −4181.32 | 1182028.55 | 13.44 | 0.31 | 259.44 | 5.73 |
| *Ice Pleurosigma sp.* | −3779.99 | 1056532.30 | 10.37 | 0.27 | 262.63 | 5.19 |
| *Temperate Navicula sp.* | −5713.27 | 1698856.24 | 77.82 | 1.17 | 245.72 | 7.58 |
| *Temperate Pleurosigma sp.* | −5322.41 | 1580474.72 | 57.68 | 0.90 | 249.74 | 8.25 |
| *Temperate Pinnularia sp.* | −6417.13 | 1846748.73 | 174.28 | 0.29 | 229.77 | 9.43 |

**Table S4.** Shape parameters and weights for distributions of ice diatom motility at various temperatures.

| $T$ (°C) | $\alpha$ | $\beta$ | $k$ | $\chi$ |
|---|---|---|---|---|
| $-7$ | 0.51 | 600.00 | 1.10 | 0.40 |
| $-7$ | 3.05 | 906.77 | 1.11 | 0.34 |
| $-5$ | 0.40 | 398.79 | 1.41 | 0.50 |
| $-5$ | 0.20 | 502.18 | 1.79 | 0.51 |
| $-3$ | 0.43 | 133.29 | 5.13 | 0.82 |
| $-3$ | 0.30 | 149.89 | 4.01 | 0.70 |
| $-1$ | 0.38 | 140.80 | 4.13 | 0.80 |
| $-1$ | 0.40 | 1187.56 | 2.15 | 0.70 |
| 0 | 0.20 | 8324.62 | 4.31 | 0.62 |
| 0 | 0.19 | 9782.45 | 4.60 | 0.60 |
| 1 | 0.25 | 989.12 | 3.50 | 0.68 |
| 1 | 0.16 | 8017.00 | 4.77 | 0.60 |
| 3 | 0.33 | 329.01 | 6.54 | 0.89 |
| 3 | 0.26 | 788.14 | 6.43 | 0.89 |
| 6 | 0.26 | 2146.14 | 5.06 | 0.75 |
| 6 | 0.40 | 762.84 | 6.18 | 0.61 |
| 12 | 0.39 | 461.42 | 8.33 | 0.76 |
| 12 | 0.31 | 4543.56 | 9.27 | 0.60 |
| 18 | 0.21 | 1988.65 | 7.70 | 0.77 |

**Supplementary Video Legend**

**Movie S1.** Discoveries of ice gliding diatoms in the Arctic Ocean.

**Movie S2.** Ice diatoms gliding in ice microchannels.

**Movie S3.** Ice and temperate diatoms gliding on glass vs. ice substrates.

**Movie S4.** A gliding ice diatom interacts with substrate at tilted angles.

**Movie S5.** Ventral flux produced by ice diatoms.

**Movie S6.** Traction forces generated by gliding ice diatoms.
